# Gut-associated bacteria invade the midgut epithelium of *Aedes aegypti* and stimulate innate immunity and suppress Zika virus infection in cells

**DOI:** 10.1101/866897

**Authors:** Shivanand Hegde, Denis Voronin, Aitor Casas-Sanchez, Miguel A. Saldaña, Eva Heinz, Alvaro Acosta-Serrano, Vsevolod L. Popov, Ashok K. Chopra, Grant L. Hughes

## Abstract

Microbiota within mosquitoes influence nutrition, immunity, fecundity, and the capacity to transmit pathogens. Despite their importance, we have a limited understanding of host-microbiota interactions, especially at the cellular level. It is evident bacterial symbionts that are localized within the midgut also infect other organs within the mosquito; however, the route these symbionts take to colonize other tissues is unknown. Here, utilizing the gentamicin protection assay, we showed that the bacterial symbionts *Cedecea* and *Serratia* have the capacity to invade and reside intracellularly within mosquito cells. Symbiotic bacteria were found within a vacuole and bacterial replication was observed in mosquito cell by transmission electron microscopy, indicating bacteria were adapted to the intracellular milieu. Using gene silencing, we determined that bacteria exploited host factors, including actin and integrin receptors, to actively invade mosquito cells. As microbiota can affect pathogens within mosquitoes, we examined the influence of intracellular symbionts on Zika virus (ZIKV) infection. Mosquito cells harbouring intracellular bacteria had significantly less ZIKV compared to uninfected cells or cells exposed to non-invasive bacteria. Intracellular bacteria were observed to substantially upregulate the Toll and IMD innate immune pathways, providing a possible mechanism mediating these anti-viral effects. Examining mono-axenically infected mosquitoes using transmission electron and fluorescent microscopy revealed that bacteria occupied an intracellular niche *in vivo*. Our results provided evidence that bacteria that associate with the midgut of mosquitoes have intracellular lifestyles which likely have implications for mosquito biology and pathogen infection. This study expands our understanding of host-microbiota interactions in mosquitoes, which is important as symbiont microbes are being exploited for vector control strategies.

## Introduction

Mosquitoes are holometabolous insects with aquatic and terrestrial life stages. Aquatic stages are continually exposed to microbes in the larval habitat while adults likely acquire microbiota from the environment after eclosion or when nectar feeding [1–3]. Additionally, environmentally acquired microbes may persist in mosquito tissues between aquatic and adult life states facilitating transstadial transmission [4–6]. It is likely that these processes contribute to the considerable variability seen in the adult microbiome [7–9]. While our understanding of genetic factors that influence host-microbe interactions and microbiome acquisition are expanding [10, 11], we still have a poor knowledge of these interactions at the cellular level. Given the importance of the microbiome on mosquito traits relevant for vectorial capacity and vector competence [12–15], understanding processes that influence microbiome homeostasis is critical for developing microbial-based control strategies [16, 17].

Bacterial microbiota often resides within several organs in mosquitoes and appears to be able to migrate between tissues. Several studies have identified bacteria in the gut of mosquitoes [9, 18–20], which have led to these microbes being commonly referred to as gut microbes, but many of these bacterial species also colonize other tissues such as the salivary gland [18, 20–23], reproductive tract [20, 22, 24], or malpighian tubules [4]. While some bacteria are unique to each tissue, several infect multiple tissues within the mosquito [20, 25], and localization in organs such as the malpighian tubules and reproductive tissues likely enables transmission between life stages and generations, respectively. Both *Asaia* and *Serratia* are transferred vertically to progeny after administered to the mosquito in a sugar meal, suggesting symbiotic bacteria have the capacity to translocate from the midgut to the germline of their host [20, 26–28]. However, mechanisms facilitating their translocation remain elusive. Infection of the entomopathogic fungus *Beavaria* of *Anopheles* mosquitoes enabled *Serratia* to escape the midgut and over replicate in the hemolymph, which was the cause of mortality to the insect [29]. In *Drosophila*, orally infected *Serratia* localized within the midgut epithelium [30]. While the epithelial infection was rare in wild type flies, *Serratia* was observed to localize intracellularly in *imd* knock-out flies, suggesting that host immunity influenced cellular localization or controlled infections [30]. Although intracellular bacterial infections have been observed in the midgut of flies [30], cellularity of gut-associated bacterial infections in mosquitoes and the mechaimism facilitating systemic infection of different tissues is largely unknown. It is plausible that an intracellular lifestyle could provide a mechanism for transstadial and vertical transmission of bacteria in mosquito vectors.

In mammalian systems, bacteria exploit their invasive capability to colonize host tissue and systemically spread within multicellular organisms [31–33]. Pathogenic bacteria like *Listeria, Salmonella, Vibrio,* and *Yersinia* invade host cells to colonize, replicate, and migrate between cells [32]. While the invasive capacity and mechanisms have been studied extensively in mammalian cells, *in vitro* investigation in mosquitoes or other arthropod vectors is lacking. In order to obtain a more complete understanding of the cellularity of bacteria associated with mosquitoes, we assessed the invasive capability of two common *Enterobacteriaceae* bacteria in mosquito cells using the gentamicin invasion assay. Using this *in vitro* assay, we characterized the invasive process, examined the mechanisms by which bacteria invade cells, and assessed the effect of intracellular bacteria on host immunity and Zika virus (ZIKV) infection. Importantly, using mono-axenically infected mosquitoes, we found that these bacteria have intracellular localization in mosquitoes. This work expands our understanding of host-microbe interactions of gut-associated symbionts in medically relevant mosquito vectors at the cellular level.

## Results and Discussion

### Symbiotic bacteria invade mosquito cells *in vitro*

Horizontally acquired bacteria are generally considered to infect the gut lumen, they are also found in other organs of mosquitoes including the salivary glands, malpighian tubules, and germline [3, 4, 6, 25, 34]. It remains unknown how these tissues become infected, but it has been proposed these organs may act as a reservoir to facilitate transstadial transmission of microbes between mosquito life stages [4, 5]. One possibility is that gut bacteria exploit their intracellular lifestyle to transition between host tissues. Therefore, we investigated the capacity of bacteria commonly found in the gut of mosquitoes to invade mosquito host cells. We isolated two bacteria within the *Enterobacteriaceae* family, *Serratia* sp. and *Cedecea neteri,* by conventional microbiological culturing approaches, and evaluated their invasive capacity using the gentamicin invasion assay [35].

While invasion assays are routinely used for pathogenic bacteria in mammalian cells, the assay is not commonly undertaken with mosquito cells. We performed the gentamicin invasion assay in Aag2 (*Aedes aegypti*) and Sua5B (*Anopheles gambiae*) cell lines comparing the invasion of *E. coli* BL21 (DE3) with or without the *Yersinia pseudotuberculosis* (*Yp*) *invasin* (*inv*) gene to invasion of these bacteria in Vero cells (Monkey Kidney cells). In mammalian systems, heterologous expression of the *Ypinv* gene facilitates invasion of *E. coli* into cell lines [36–38]. Similar to mammalian systems, we found that *E. coli* expressing the *Ypinv* gene had significantly increased invasion in Aag2 cells compared to the non-invasive *E. coli* control (Fig. S1, Unpaired t test, p < 0.05), while no statistical difference was seen in the Sua5B cell line, likely due to high variability among replicates (p > 0.05, Unpaired t test). As expected*, E. coli* expressing the *inv* gene invaded at significantly higher rates in Vero cells compared to non-invasive *E. coli* (p < 0.05, Unpaired t test). While several insect cell lines are naturally phagocytic [39, 40], our data suggested bacteria were actively invading Aag2 cells, and as such, we conducted the majority of our experiments with this cell line. Next, we completed the gentamicin invasion assay with the two gut-associated bacteria from mosquitoes, *C. neteri* and *Serratia* sp., and used *E. coli* with and without the *Ypinv* gene as the positive and negative controls, respectively. The native symbionts exhibited significantly higher rates of invasion compared to the *E. coli* expressing *Ypinv* (ANOVA with Tukey’s multiple comparision test, p < 0.05) or wildtype *E. coli* (Fig. 1A, ANOVA with Tukey’s multiple comparision test, p < 0.01) indicating native gut-associated microbes have the capacity to invade insect cells and are more adept at this process compared to non-native *E. coli* expressing mammalian invasive factors.

**Figure 1.**
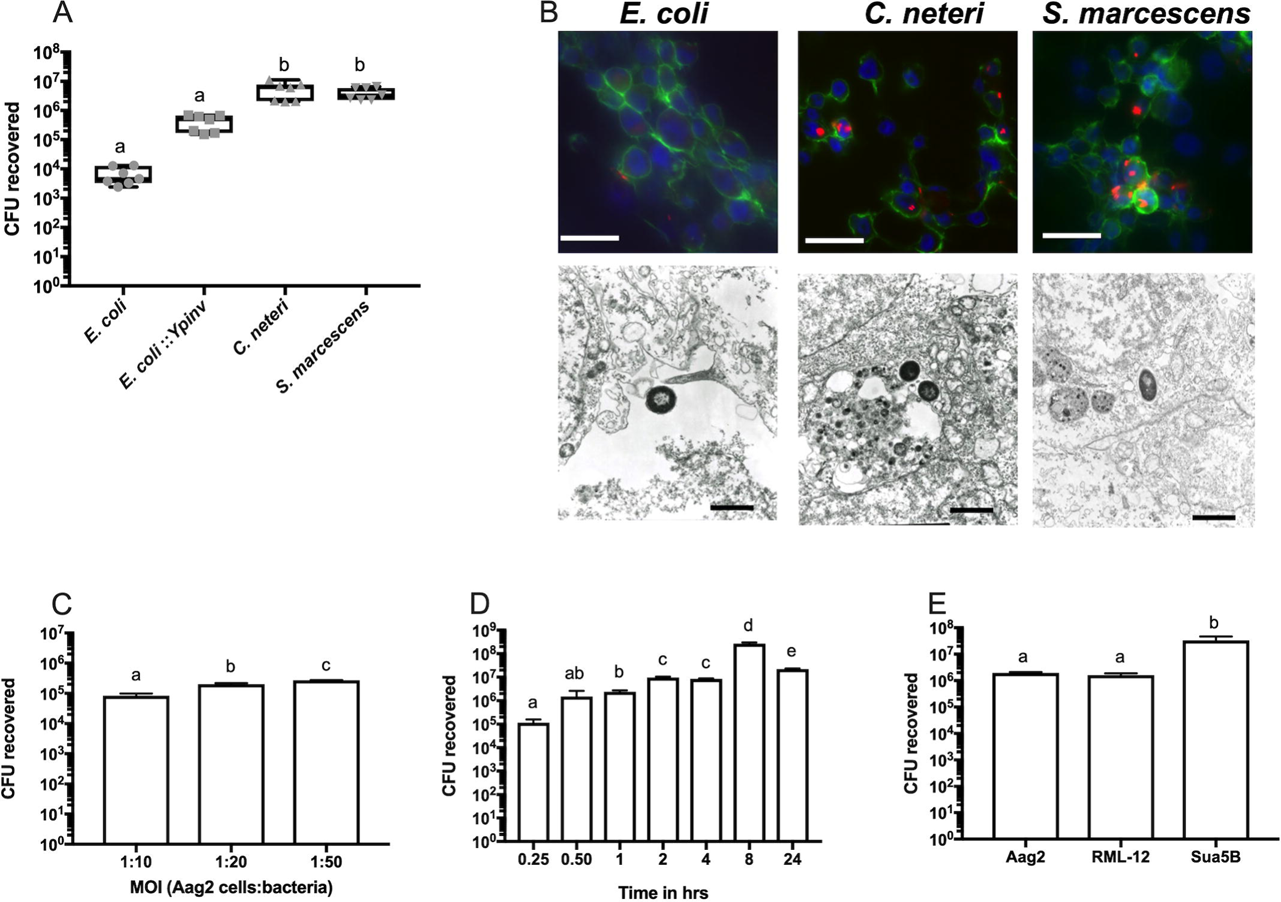
Invasion of symbiotic bacteria into mosquito cells. The gentamicin invasion assay was used to examine the invasive capacity of symbiotic *Enterobacteriaceae* bacteria isolated from *Aedes* mosquitoes (A). Non-invasive *E. coli* was used as negative control. *E. coli* expressing the *Yersinia inv (Ypinv)* gene was used as a positive control. The assay was repeated twice. Flourscent and transmission electron microscopy was used to visual intracellular bacteria (B). Bacteria expressed mCherry fluorescent protein (red), actin filaments were stained with Phalloidin (green) and DNA with DAPI (blue). Arrowheads in the TEM images indicate vacuoles containing bacteria. Scale bar is 500 nm. Density (C) and time dependent (D) invasion of *C. neteri* in Aag2 cells. The density dependant invasion assay (C) was replicated twice. The time dependant invasion assay was done at host cell: bacterial density of 1:10 (N=4). *C. neteri* invasion in *Aedes aegypti (*Aag2 and RML-12) *and Anopheles gambiae* (Sua5B) (E). The assay was repeated twice. Letters indicate significant differences (p < 0.05) determined by a One-Way ANOVA with a Tukey’s multiple comparison test.

To further confirm the results from the gentamicin invasion assay, fluorescent and transmission electron microscopy (TEM) were performed on cells after invasion. In order to observe the invaded bacteria in cells using fluorescent microscopy, bacteria were transformed with a plasmid that expressed the mCherry fluorescent protein [41]. Similar to our quantitative results from the invasion assay, we observed a greater number of intracellular bacteria in the *Serratia* and *Cedecea* treatments compared to the *E. coli* negative control (Fig. 1B and Fig. S2). TEM images confirmed both symbionts isolated from mosquitoes were intracellular, and that bacteria were inside a vacuole (Fig. 1B, black arrowhead). *E. coli* did not invade cells and was found exclusively extracellularly. Bacterial encapsulation within a vacuole is a typical signature of invading bacteria in mammalian systems [42] as well as obligatory intracellular bacteria of insects such as *Wolbachia* [43]. Taken together, it is evident that both symbionts isolated from the mosquitoes can invade the host cells *in vitro*.

Next, we characterized the invasion process of *Cedecea* examining how the multiplicity of infection (MOI) and incubation time influenced invasion. We noted a linear increase in the number of intracellular *Cedecea* with increasing multiplicity of infection (MOI) (Fig. 1C, ANOVA with Tukey’s multiple comparision test, p < 0.05). We then varied the invasion time and observed bacterial invasion as early as 15 minutes post infection and invasion increased until 8 hr post infection (Fig. 1D, ANOVA with Tukey’s multiple comparision test, p < 0.05). We also examined the invasive ability of *C. neteri* in different mosquito cells lines. The invasion of this bacterium was similar in both *Ae. aegypti* cell lines Aag2 and RML-12 (Fig. 1E, Tukey’s multiple comparision test, p < 0.05); however, a greater number of intracellular bacteria were seen in the Sua5B cells.

### *In vitro* intracellular replication and egression of *Cedecea neteri*

While undertaking TEM, we captured an image of intracellular replication of *C. neteri* within mosquito cells (Fig. 2A). Given that bacteria were seen to replicate in the intracellular environment, we attempted to culture these bacteria in Aag2 cells in a similar manner to *in vitro* propagation of other intracellular bacteria such as *Wolbachia* [44]. However, our culturing attempts were unsuccessful as the cell culture media became contaminated with the innoculated *C. neteri,* despite the extracellular bacteria, which had not invaded, being killed by gentamicin treatment. We hypothesized that intracellular *Cedecea* were egressing from the cells and replicating within the cell culture media.

**Figure 2.**
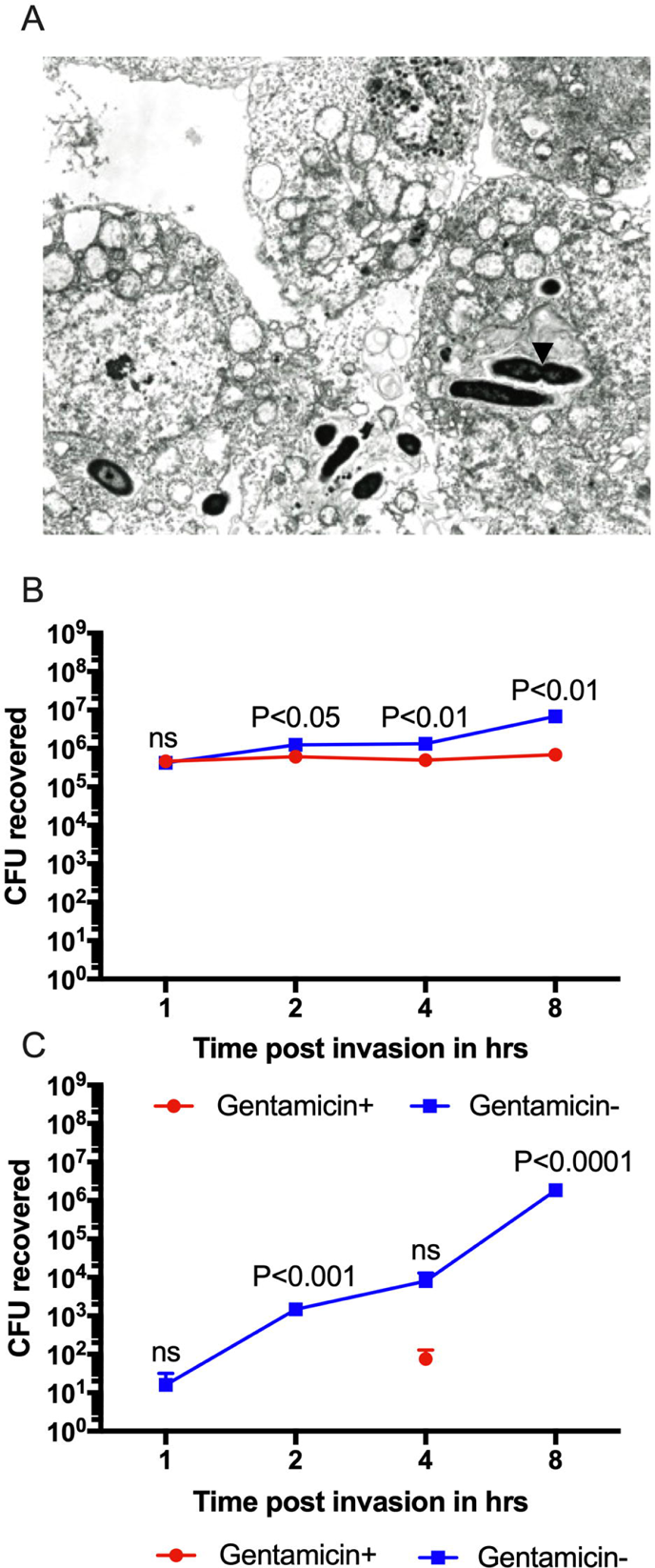
Intracellular replication and egression of *C. neteri* in Aag2 cells. TEM of Aag2 cells containing invaded *C. neteri* replicating inside Aag2 cell (A). Arrow indicates the dividing bacterial cell. Bacterial titer in cells (B) or in the cell culture media (C) in the presence and absence of gentamicin. The significance between the gentamicin and non-treated samples at different time post invasion was analyzed by Unpaired t test. Five replicates were used at each time point.

We therefore undertook experiments to quantify bacterial egression from the cells. After allowing *C. neteri* to invade, Aag2 cells were incubated with or without gentamicin and intracellular and extracellular bacteria were quantified over time. Within the cell, bacterial numbers remained constant in the presence of gentamicin while in the absence of antibiotic, there was an an approximate 10-fold increase at eight hours post infection (Fig. 2B, Unpaired t test, p<0.05). In the cell culture media, we observed a precipitous increase in bacteria in the absence of antibiotic and recovered little to no viable bacteria when antibiotics were included in the media (Fig. 2C, Unpaired t test, p<0.05). These data indicated that *C. neteri* was egressing from the cells, replicating in the cell culture media in the absence of antibiotic and then re-invading mosquito cells, which accounted for the significantly higher titer of intracellular *Cedecea* in the non-antibiotic treated cells at 8 hours post infection. We found few changes in the total number of mosquito cells in gentamicin treated or untreated cells, although there was a subtle but significant reduction in cell number after 8 hours in the treatment without antibiotics (Fig. S3, Unpaired t test, p<0.0001). However, overall, these data indicated that bacterial invasion and egression were not overly detrimental to the host cells.

### Host actin and integrin are important for *Cedecea neteri* invasion

The lack of damage to host cells indicates non-lysis mediated exit of bacteria from host cells. While this would be expected from a mutualistic or commensal gut-associated bacterium, even some pathogens such as *Mycobacteria*, *Shigella*, and *Chlamydia* use protrusions and non-lytic exocytosis to exit the host cells without lysis of host cells [45–48]. The former method involves membrane extensions containing bacteria mediated by actin polymerization and ultimately these protruded structures are engulfed by neighboring cells resulting in the transfer of content to adjoining cell [49]. We therefore hypothesized that *C. neteri* may use actin-based motility as a mechanism to invade and exit cells.

To determine the role of the actin cytoskeleton in invasion of bacteria into mosquito cells, we inhibited the polymerization of actin filaments using cytochalasin D [50]. We observed a 3-fold reduction in invasion of *Cedecea* in the presence of cytochalasin D (Fig. 3A, ANOVA with Tukey’s multiple comparision test, p< 0.001). In contrast, there was no change in intracellular bacteria when cells were treated with SP600125, which inhibits phagocytosis in mosquito cells [51]. These data suggested that the actin cytoskeleton is co-opted by *C. neteri* to gain access to the intracellular milieu, and that phagocytosis played a minimal role in the invasion of bacteria. Similar processes have been observed in other bacteria-host systems. For example, obligatory intracellular bacteria such as *Rickettsia*, *Chlamydia*, and *Ehrlichia* hijack the host cell cytoskeletal and surface proteins to invade, survive and spread within cells [52–54]

**Figure 3.**
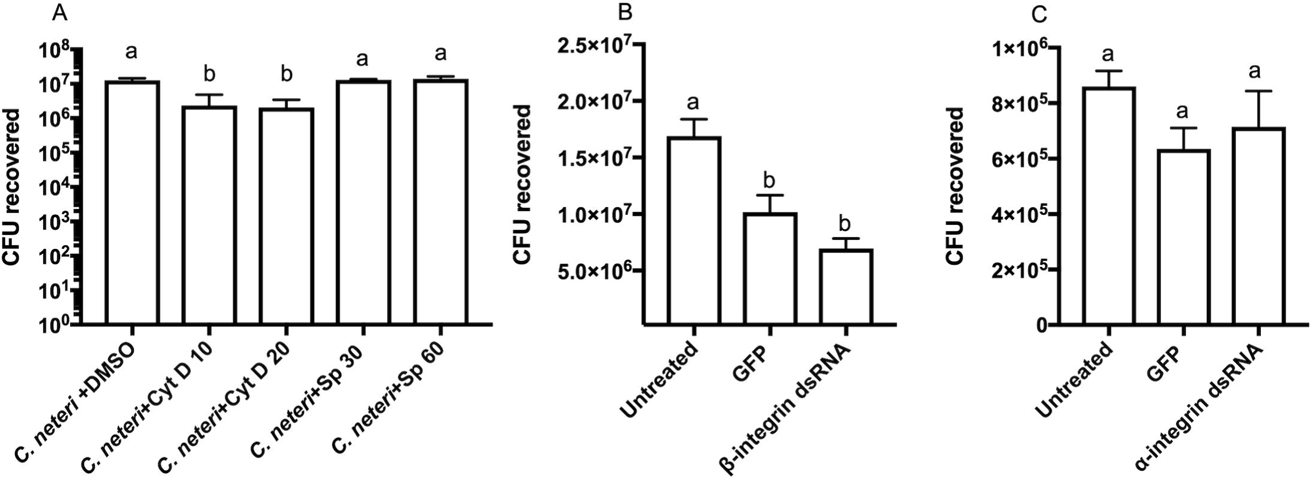
**The role of host cytoskeletal proteins and receptors in the *C. neteri* invasion**. Invasion of *C. neteri* in the presence of inhibotors of actin polymerisation (Cyt D) and phagocytosis (SP600125 [Sp]) (A). DMSO was used as control to assess its cytotoxic effect on the cells. *C. neteri* invasion after silencing of the beta-(B) and alpha-(C) integrins. The expeirments were repeated twice. Letters indicate significant differences (p < 0.05) determined by a One-Way ANOVA with a Tukey’s multiple comparison test.

We then examined whether host receptors facilitate the bacterial entry into mosquito cells. In mosquitoes, integrins are involved in the engulfment of *E. coli* and malaria parasites [55], while pathogenic bacteria of humans also exploit these receptors to invade mammalian host cells [56–58]. Using RNAi, we silenced the alpha and beta subunit of the integrin receptor and challenged cells with *C. neteri*. After confirming gene silencing (Fig. S4 A and B), we found significantly fewer intracellular bacteria after knocking down the beta-integrin (Fig. 3B, ANOVA with Tukey’s multiple comparision test, p < 0.05), but no differences in the rate of *Cedecea* invasion when the alpha-integrin gene was silenced (Fig. 3C, ANOVA with Tukey’s multiple comparision test, p > 0.05). These results indicated that symbiotic *C. neteri* utilized actin filaments and the beta-integrin receptor to gain entry into the host cells.

### Intracellular *Cedecea* reduces ZIKV replication in mosquito cells

Midgut-associated bacteria can affect pathogens transmitted by mosquitoes by direct or indirect interactions [59–61]. Therefore, we examined how intracellular *C. neteri* affected viral infection. The symbiont significantly reduced ZIKV loads in cell lines compared to uninfected controls at both two (Fig. 4A, Unpaired t test, p < 0.05) and four (Fig. 4B, Unpaired t test, p < 0.01) days post virus infection (dpvi) (Fig 4A and 4B). Similar to *Cedecea*, intracellular *Serratia* also significantly reduced ZIKV density by four logs compared to the uninfected cells at four dpvi (Fig. 4C, ANOVA with Tukey’s multiple comparison test, p < 0.05). To determine how the density of bacteria influenced viral infection, we infected cells at increasing bacterial MOIs before inoculating with virus. At lower MOIs (1:1 and 1:2), *Cedecea* significantly reduced ZIKV compared to the *E. coli* (MOI 1:1, Unpaired t test, p <0.01, MOI 1:2, Unpaired t test, p <0.0001). However, at higher MOIs, we noted that both *Cedecea* and *E. coli* reduced ZIKV compared to the uninfected control. The complete blockage of ZIKV at the higher MOIs suggested that even non-invasive bacteria can overwhelm viral infection, likely by induction of the immune effector molecules that are antagonistic to viral infection. Taken together, our results suggested that members of *Enterobacteriaceae* that commonly infect mosquitoes have the capacity to interfere with viral pathogens when they are intracellular.

**Figure 4.**
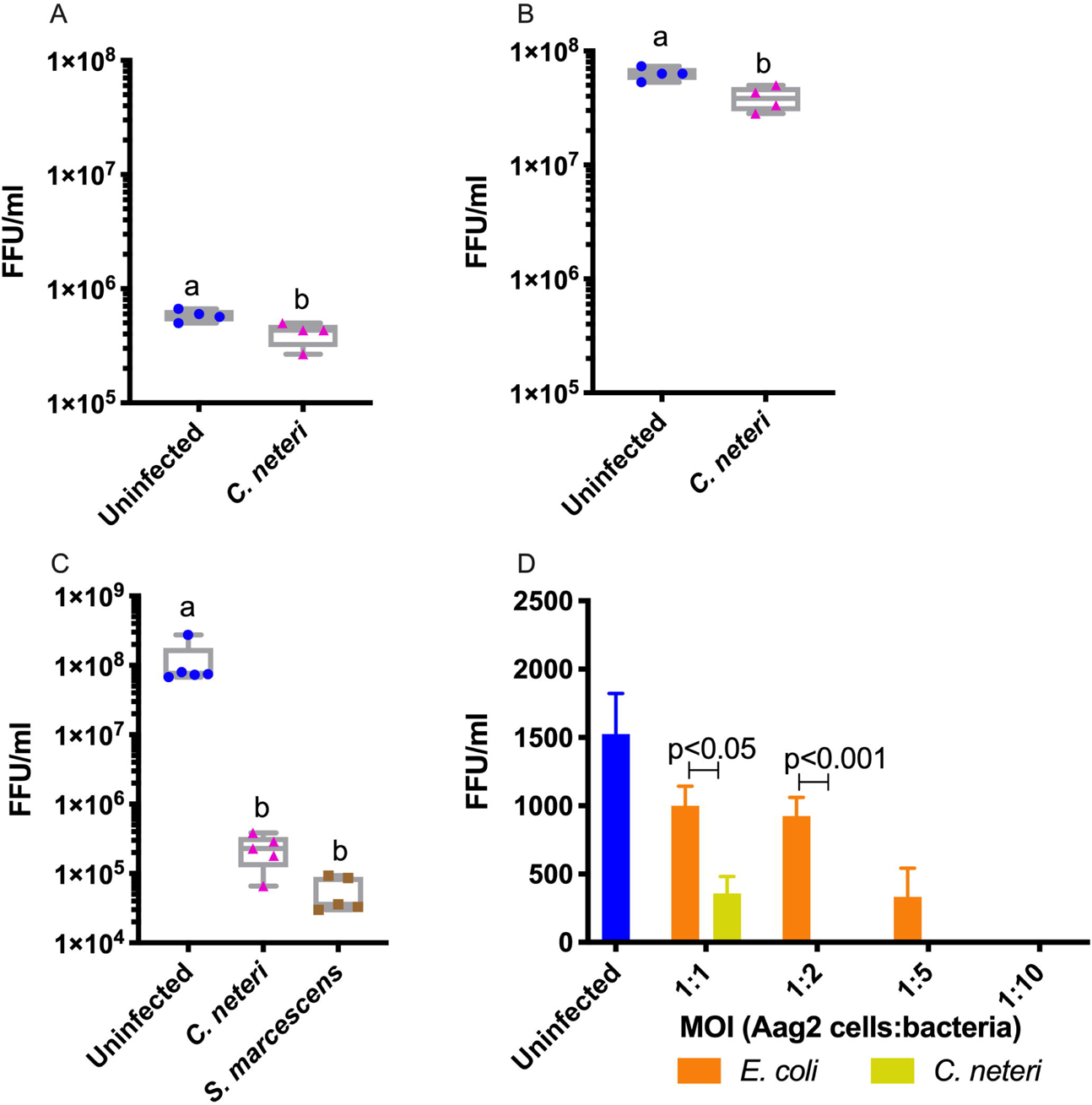
**Intracellular bacteria reduces ZIKV titer in *Ae. aegypti* cells**. ZIKV infection at 2 (A) and 4 (B) days post invasion compared to an uninfected control. ZIKV infection in *C. neteri* or *Serratia* sp. infected cells (C). The effect of bacterial density on ZIKV infection (D). *C. neteri* and *E. coli* were inoculated onto cells using the gentamicin invasion assay at increasing MOIs. For the *C. neteri* 1:2, 1:5, and 1:10 and *E. coli* 1:10 treatments, no ZIKV was recovered from cells. For A, B and D significance was determind using unpaired t-test, while for C, significance was calculated by one-way ANNOVA with Tukey’s multiple comparison test.

### *Cedecea* invasion stimulates mosquito immunity

There is a complex interplay between the host innate immune system and microbiota which maintains microbiome homeostasis [16, 62, 63]. However, invading arboviral pathogens are also susceptible to these immune pathways [64, 65] thereby providing an indirect mechanism by which microbiota can interfere with pathogens. We therefore examined the immune response of mosquito cells challenged with *Cedecea* or *E. coli* comparing these responses to uninfected cells. We quantified the transcription factors (*rRel1*, *rRel2* and *Stat*) and negative regulators (*Cactus*, *Caspar*, and *PIAS*) of the Toll, IMD and Jak/Stat immune pathways as well as downstream effector molecules (*gambicin*, *definsin* and *cecropin*). We found the NF-κB transcription factor *Rel2* was significantly upregulated by *Cedecea* compared to both the *E. coli* (ANOVA with Tukey’s multiple comparison test, p < 0.05) and the uninfected control (ANOVA with Tukey’s multiple comparison test, p < 0.01), while a significant difference was only observed for *Rel1* when the *Cedecea* treatment was compared to the uninfected control (ANOVA with Tukey’s multiple comparison test, p < 0.05; Fig 5A). The negative regulator of the Toll pathway, *Cactus*, was significantly upregulated compared to both the *E. coli* (Fig. 5B, ANOVA with Tukey’s multiple comparison test, P < 0.05) and uninfected control (ANOVA with Tukey’s multiple comparison test, p < 0.01), while no changes were seen for *Caspar*, the negative regulator of the IMD pathway (Fig. 5B, ANOVA with Tukey’s multiple comparison test, p>0.05,). Similarly, no significant changes were observed for genes in the Jak/Stat pathway. Taken together, these data suggested that the Toll and IMD pathways were induced by invasion of *Cedecea* into mosquito cells. This is consistent with previous observations which demonstrated interplay between native microbiota and mosquito innate immune pathways [61, 66–68]. We observed dramatic modulation of effector molecules with *Defensin*, *Cecropin* and *Gambicin,* all significantly enhanced by *Cedecea*. Strikingly, *Cecropin* and *Defensin* expression was nearly 1000-fold higher (Fig. 5C, Tukey’s multiple comparison test, p < 0.0001) whereas *Gambicin* (Fig. 5C, Tukey’s multiple comparison test, p <0.001) was elevated 100-fold in cells inoculated with *Cedecea* compared to the non-infected control. In mosquitoes, these downstream effector molecules are co-regulated by the Toll and IMD pathways [66], which could explain their prolific enhancement given that intracellular *Cedecea* stimulated both pathways. As arboviral pathogens also interact with innate immune pathways, we examined gene expression in cells when co-infected with ZIKV and *C. neteri* focusing on the NF-κB transcription factors and negative regulators of the Toll and IMD pathways. Patterns of gene expression were similar to the ZIKV uninfected cells with the exception of *Cactus*, where no significant differences were seen across bacterial treatments (Fig. 5D and E), suggesting that ZIKV was stimulating the Toll pathway as the negative regulator was depleted when comparing to ZIKV uninfected cells.

**Figure 5.**
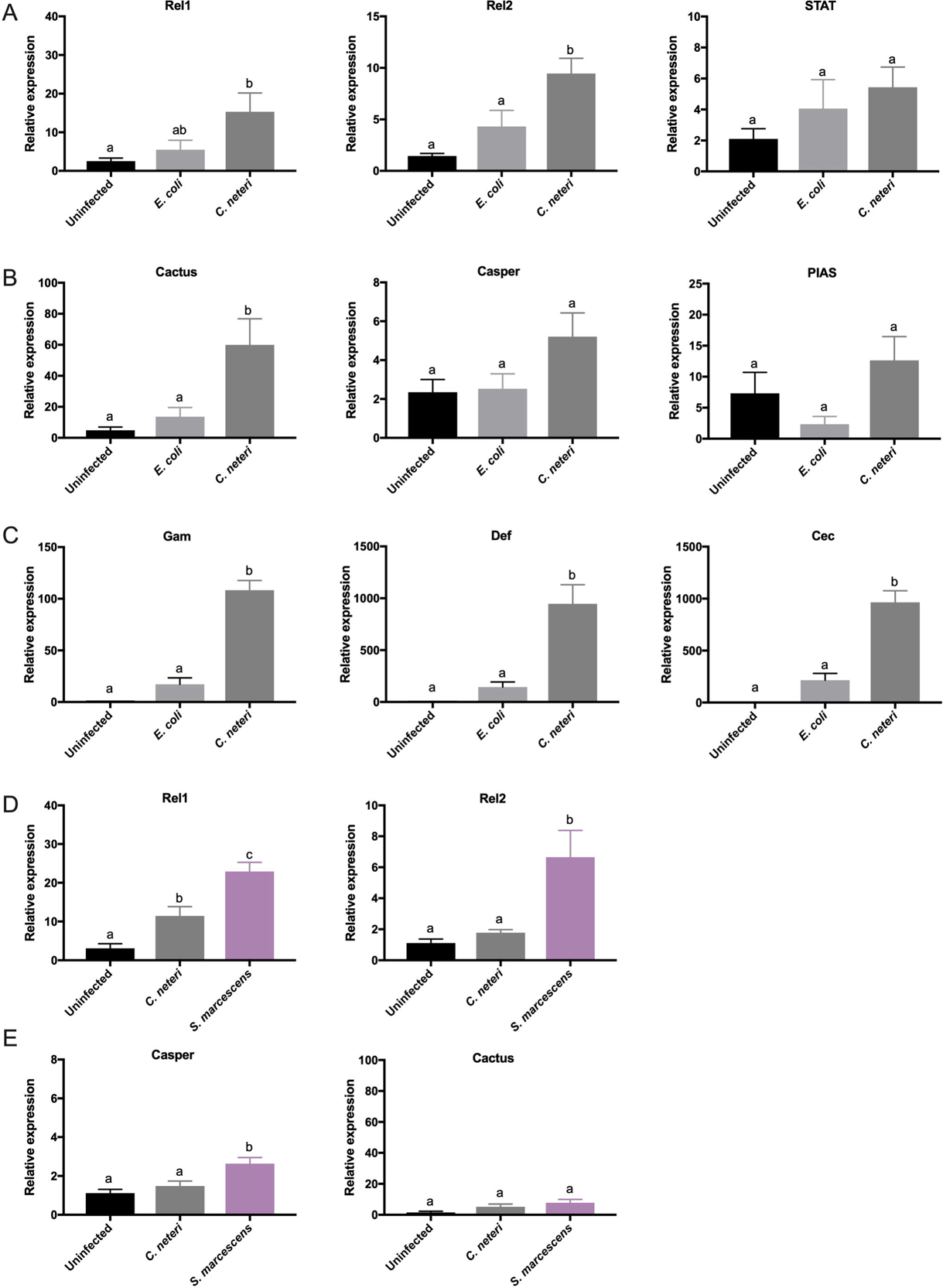
Intracellular *C. neteri* upregulates mosquito Toll and IMD innate immune pathways. Gene expression analysis of the NF-κB transcriptional activators (A) and the negative regulators (B) of the Toll, IMD and JAK-STAT pathways as well as downstream effector molecules (C). Gene expression was measured 24 hr post *C. neteri* invasion in Aag2 cells. Gene expression of the NF-κB transcriptional activators (D) and the negative regulators (E) in cells co-infected with *C. neteri* or *Serratia* sp. and ZIKV. 24 hours post bacterial infection cells were infected with ZIKV. Samples were collected 4 days post ZIKV infection for qPCR analysis. The experiment was repeated twice. Letters indicate significant differences (p < 0.05) determined by a One-Way ANOVA with a Tukey’s multiple comparison test.

### Intracellular *Enterobacteriaceae* within the *Aedes* gut epithelium

To determine the capacity of *C. neteri* and *Serratia* sp. to invade host cells *in vivo,* we reared *Ae. aegypti* mosquitoes mono-axenically with either symbiont and analyzed tissues from larvae and adults by TEM and Confocal Laser Scanning Microscopy (CLSM). While it was evident there was an accumulation of extracellular *Serratia* in the lumen of the larval gut (Fig. 6A-B), we also identified bacteria that were associated with the microvilli. Specifically, we found examples of *Serratia* in the process of transitioning to or from the midgut epithelial cells. We appreciate our results cannot conclusively determine if *Serratia* was in the process of invading or egressing from cells, but regardless, it suggested that the bacterium had been or was soon to be intracellular. Analysis of gut tissue isolated from adult *Ae. aegypti* mosquitoes infected with *C. neteri* revealed the presence of bacteria in the cytosol of epithelial cells (Fig. 6B). Closer inspection of these images revealed the bacterium was localized within a vacuole (Fig. 6B, yellow insert), which is a typical signature of intracellular bacteria. Here, *C. neteri* may be exiting the membrane (Fig. 6B, inserts, white arrow), suggesting these bacteria can egress from the membrane bound compartment which could facilitate their replication and spreading. Egression and re-entry mechanisms are used by several pathogenic bacteria like *Listeria monocytogenes* and *Shigella flexneri* to escape the vacuoles to replicative niches [69]. We also confirmed the intracellular localization of *C. neteri* in adult infected guts by CLSM. The orthogonal views of the 3D-reconstructed tissues locate bacteria on the cells as well as inside cells of the posterior gut, demonstrated by the co-localization of actin staining with mCherry signal from bacteria (Fig. 6C, Fig. S5, Supplementary video 1). We also found bacteria inside the cells of the Malpighian tubules (Fig. 6C, Fig. S5, Supplementary video 2). Altogether, TEM and CLSM results clearly show bacteria residing inside the host cells *in vivo*.

**Figure 6.**
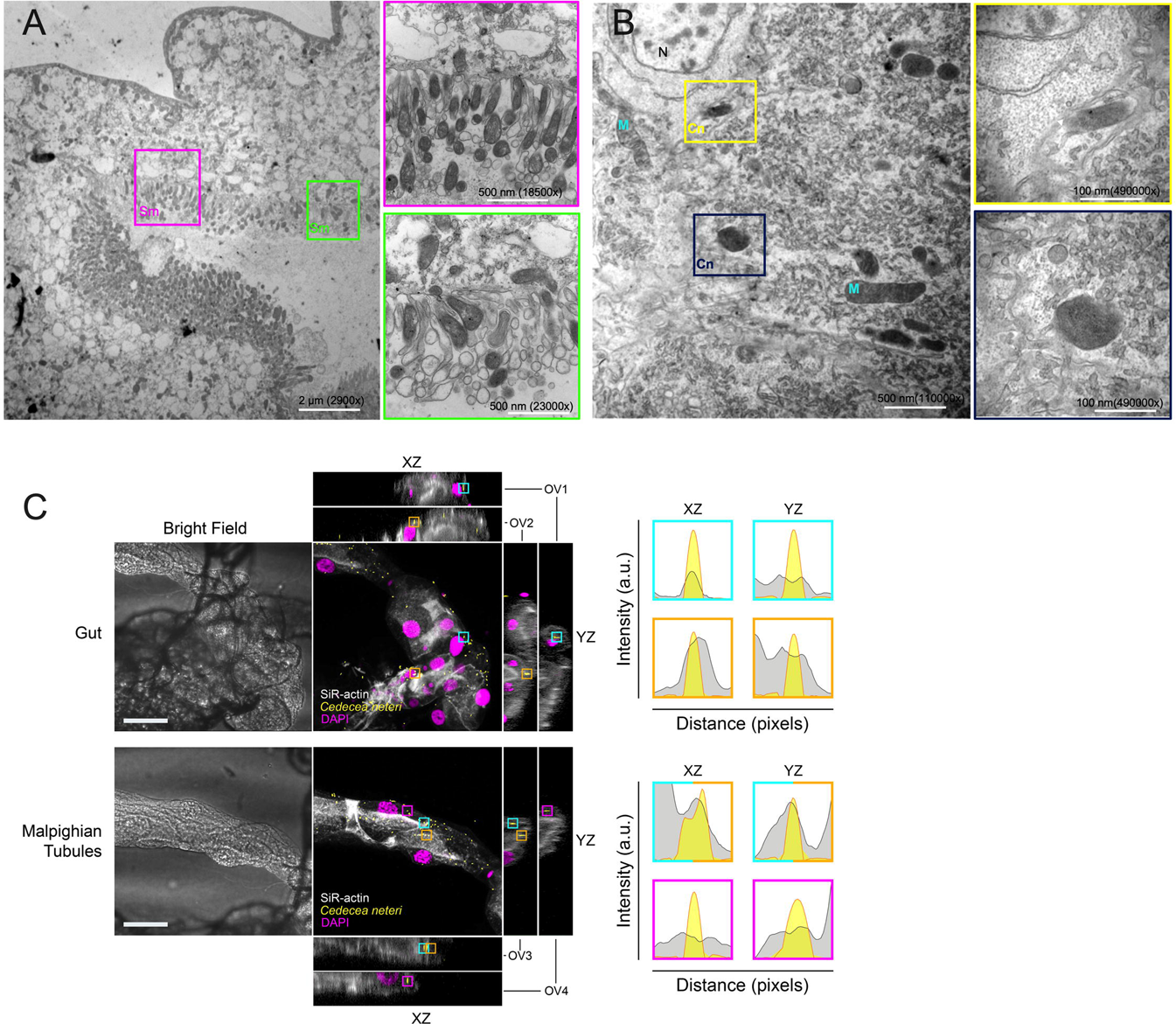
Intracellular localization of *Cedecea* and *Serratia* in mosquito tissues. TEM micrographs of *Serratia* sp. (Sm) accumulated in the gut lumen and associated with the microvilli of the gut epithelium in mono-axenically infected *Ae. aegypti* adults (A). Magnified image of bacteria attaching to microvilli (MV) (B), and bacteria in the process of entering or exiting the gut epithelia (purple and green insert). (B) Intracellular *C. neteri* (Cn) in the larval gut mono-axenically infected *Ae. aegypti*. Mitochondria (M) and nucleus (N). Yellow and blue inserts show larger view of bacteria from B. CLSM evidence of the intracellular localization of *C. neteri* in the adult mosquito gut and Malpighian tubules (C). Bright field (left) and maximum intensity projection (right) of tissues 3D-reconstructed from a series of Z-stacks merging mosquito actin (white), mCherry-expressing *C. neteri* (yellow) and DAPI-stained DNA (magenta). Two representative XZ and YZ orthogonal views (OV1-4) of the stacks are shown for each tissue on the sides, and the identity of intracellular bacteria examples is noted with colored squares. On the right, the plots coloured according to the identity of the corresponding bacterium, show the co-localization of the actin signal (gray) with the mCherry bacteria (yellow). Scale bars are 50 μm.

Our data show that *Enterobacteriaceae* that commonly infect the gut of mosquitoes have the capacity to invade mosquito cells *in vitro* and *in vivo*. To gain access and persist in these cells, bacteria need to overcome the host immune response and the peritrophic matrix (PM). The PM, which acts as a physical barrier that separates epithelial cells from the gut lumen, is expressed constitutively in larvae and after a blood meal in adults. In a range of arthropods, genes associated with the PM are induced by bacteria [70] and in turn, the PM plays a pivotal role maintaining gut microbiome homeostasis, either by protecting bacteria from the innate immune response or restoring bacterial composition and adundance post blood meal [71]. While another study has identified bacteria associating with the epithelium in *Anopheles* mosquitoes [72] our finding of intracellular bacteria residing within the midgut epithelium of larvae and adults indicates the PM is not completely effective at inhibiting microbiota or that bacteria invade these cells before the PM has established. Alternatively, bacteria may produce enzymes that degrade the PM in a similar fashion to malaria parasites that express chitenases [73].

In *Drosophila*, ingested *Serratia* (Db11) invaded the midgut epithlieum in flies with an impaired IMD pathway, but not wild type flies. However, this infection reduces survival [30]. Similarly, fungal infection of *Anopheles* mosquitoes enables gut bacteria to translocate to the hemolymph leading to systemic infection [29]. Similar to the observations in *Drosophila* [30], we saw intracellular bacteria infrequently in the mosquito gut, suggesting there were intrinsic factors limiting the systemic infection. Innate immunity may be respsonsible for maintaining homeostatsis, which would be consistent with our gene expression data, or alternatively, these mutualistic bacteria may exploit similar molecular processes as their pathogenic counterparts to overcome host immune pathways [74]. The intracellular lifestyle of bacteria and their ability to egress from cells likely facilitates microbial persistence in these holometabolous insects. Our finding of intracellular bacteria in the malpighian tubules further supports this theory as bacteria residing within this tissue are known to be transstadially transmitted [4].

## Conclusions

In conclusion, we have shown through various *in vitro* and *in vivo* data that symbiotic *Enterobacteriaceae* can invade and replicate intracellularly in mosquito cells. Bacterial invasion is mediated by host actin filaments and beta-integrin receptors. Intracellular bacteria dramatically upregulate host IMD and Toll immune pathways and substantially reduce ZIKV density in cells. These data enhance our understanding of host-microbe interactions in mosquitoes and point to a possible mechanism by which bacteria, which are commonly associated with the midgut, could infect other tissues within mosquitoes.

## Material and Methods

### Ethics statement

ZIKV, which was originally isolated from an *Ae. aegypti* mosquito (Chiapas State, Mexico), was obtained from the World Reference Center for Emerging Viruses and Arboviruses at the University of Texas Medical Branch (Galveston, TX). Experimental work with the virus was approved by the University of Texas Medical Branch Institutional Biosafety Committee (reference number 2016055).

### Isolation of bacteria from mosquitoes

Lab reared *Ae. albopictus* mosquitoes were collected and surface sterilized before homogenized in 500 μl of 1X PBS. Serial dilution of homogenates was plated on LB agar plate to obtain isolated colonies. The single colonies were picked, grown in LB medium before isolating genomic DNA. The 16S rDNA PCR was performed as described previously [75] and the PCR product was Sanger sequenced to identify the bacterial species. To further classify the gut-associated bacteria we completed multilocus sequence typing (MLST) as described previously [11, 76]. The MLST sequences were aligned, concatenated and a maximum likelihood tree under the LG model and rapid bootstrap was constructed using Seaview [77] (Fig. S6). The sequences have been deposited under accession numbers XXXX.

### Bacterial growth and cell culture

Two bacterial isolates were grown in LB medium at 37°C. The overnight culture was appropriately diluted in Schneider’s media (Gibco) to obtain the MOI of 10 before the infection. The mosquito cell lines were maintained in their respective medium at 28 °C. The *Ae. aegypti* cell line Aag2 [78] and Sua5B cells were maintained in *Drosophila* Schneider’s medium (Gibco) supplemented with 20% FBS (Denville Scientific) and 1% penicillin/streptomycin (P/S; 100 Units/mL and 100 μg/mL respectively), RML-12 cells were maintained in Leibovitz’ (L15) medium (Gibco) containing 20% FBS and 10% tryptose phosphate broth. Vero cells (CCL-81) were purchased from the American Type Culture Collection (Bethesda, MD, USA) and maintained in DMEM supplemented with 5% FBS and 1% P/S (100 Units/mL and 100 μg/mL respectively) at 37 °C with 5% CO_2_.

### Gentamicin invasion assay

The gentamicin invasion assay was performed as described elsewhere with minor alterations [33]. Aag2 cells were seeded at the density of 1×10^5^/well in 24-well plate 48h prior to infection. On the day of infection, cells were washed in Schneider’s media (Gibco) and infected with 500 μl of bacterial suspension. After incubating for 1h at 28 °C, bacteria were removed, and cells were washed once with Schneider’s medium and incubated with 200 μg/ml gentamicin for additional 1h to kill extracellular bacteria. The invaded bacteria were recovered after washing the cells twice with Schneider’s media and lysing them in 500 μl of 1X PBS containing 0.05% Triton X-100.

### Fluorescence and Transmission electron microscopy

In order to assess the invasion of symbionts fluorescent microscopy and TEM was performed on the Aag2 cells after allowing bacteria to invade. The bacteria were transformed with mCherry expressing plasmid pRAM18dRGA-mCherry, which is a modified version of pRAM18dRGA[MCS] [41]. Aag2 cells were fixed with 1% PFA (Electron Microscopy Sciences, Hartfield, PA) for 30 min and permeabilized in 1X PBS+0.01 % Triton X-100 (Fischer Scientific) for 20 min following staining with Atto 488 Phalloidin (Sigma) as per manufacturers recommendations. The cell nuclei were stained with DAPI after washing the slides in 1X PBS. The slides were stored in Prolong-Antifade (Invitrogen). The samples were observed using the Revolve-FL epifluorescence microscope (ECHOLAB). For TEM, insect cells were fixed in fixative (2.5% formaldehyde, 0.1% glutaraldehyde, 0.03% picric acid, 0.03% CaCl_2_ and 0.05 M cacodylate buffer at pH 7.3) and post fixed in 1% osmium tetroxide for 1 h, stained *en bloc* in 2% aqueous uranyl acetate at 60 °C for 20 min, dehydrated in a graded series of ethanol concentrations, and embedded in epoxy resin, Poly/Bed 812. Ultrathin sections were cut on a Leica EM UC7 (Leica Microsystems, Buffalo Grove, IL), placed on Formvar-carbon copper 200 mesh grids, stained with lead citrate and examined in a Philips (FEI) CM-100 electron microscope at 60 kV. To assess the *in vivo* invasion in mosquito larvae and adults, guts were dissected after surface sterilization in 1X PBS and then the tissue was fixed in fixative (2.5% glutaraldehyde and 2% paraformaldehyde buffered with 0.1 M sodium cacodylate) for 2 hours and post fixed in 1% osmium tetroxide for 1 h at room temperature. Then samples were dehydrated in a graded series of ethanol concentrations, and embedded in epoxy resin, Epon 812. The sections were prepared as described above and imaged in a Tecnai Spirit (FEI) transmission electron microscope at 80 kV. For Confocal Laser Scanning Microscopy, tissue samples were fixed in 1% PFA in 1X PBS for 30 min, then permebalized with 0.01% Triton X-100 in 1X PBS for 20 min before staining with SiR-actin Kit (Spirochrome AG, Switzerland) for 1 hour and DAPI (Applied Biosystems) for 15 min. Then tissue samples were embedded in 1% low-melting agarose with SlowFade Diamond mounting solution (Molecular Probes). Samples were imaged and 3D-reconstructed (1.3 mm sections) using a Zeiss LSM-800 and were analysed in Zen 3.0 (Zeiss) and Fiji (ImageJ).

### Intracellular replication of *Cedecea neteri*

To assess the replication of bacteria inside the host cells as well as in the medium, the Aag2 cells innoculated with *C. neteri* were incubated with or without gentamicin for 8h at 28 °C. Every two hours, the supernatant was collected and serial dilutions were plated on LB agar plate to enumerate the bacterial quantity in the medium. The cells were washed two times with Schneider’s medium before plating on agar plate.

### Host cytoskeleton and Janus Kinase in *Cedecea neteri* invasion

The gentamicin invasion assay was performed in the presence of actin and Janus kinase (JAK) inhibitors. The assay was performed by pre-incubating Aag2 cells in presence of 10 or 20 μg/ml of Cytochalasin D (Sigma) and 30 or 60 μg/ml of Sp600125 (Sigma) for 1 hr. The gentamicin invasion assay was performed as described above with the addition of each specific drug. A 60 μg/ml of DMSO treatment was used as a control.

### RNAi mediated integrin gene silencing in Aag2 cells

In order to assess the role of host integrin receptors in the invasion of *C. neteri*, the integrin alpha and beta receptors were depleted using RNAi. dsRNA was designed for AAEL001829 and AAEL014660 using E-RNAi [79] and amplified using primers with flanking T7 promoter sequence using *Ae. aegypti* cDNA as a template. dsRNA was synthesized using the T7 megascript kit (Ambion). The primers are listed in the Table S1. dsDNA against GFP was used as control. Aag2 cells were transfected with 0.5 μg of each dsRNA using Lipofectamine™ RNAiMAX (Life Technologies) 48hrs prior to bacterial infection and the gentamicin invasion assay.

### RT-qPCR analysis

Total RNA was isolated from Aag2 cells and reverse transcribed using the amfiRivert cDNA synthesis Platinum master mix (GenDepot, Barker, TX, USA) containing a mixture of oligo(dT)18 and random hexamers. Real-time quantification was performed in a StepOnePlus instrument (Applied Biosystems, Foster City, CA) in a 10 μl reaction mixture containing 10-20 ng cDNA template, 1X PowerUp SYBR green master mix (Applied Biosystems), and 1 μM (each) primer. The analysis was performed using the threshold cycle (ΔΔCT) (Livak) method [80]. Four independent biological replicates were conducted, and all PCRs were performed in duplicate. In order to assess the expression of innate immune genes, the invasion assay was performed as described earlier and post 24-hr invasion cells were harvested to isolate RNA, followed by cDNA synthesis and RT-qPCR for specific genes. The ribosomal protein S7 gene [81] was used for normalization of cDNA templates. Primer sequences are listed in Table S1.

### *In vitro* vector competence of ZIKV in Aag2 cells

The assay was performed in order to assess the how intracellular bacteria modulate ZIKV infection *in vitro*. After the gentamicin invasion assay with *C. neteri* at an MOI of 1:1, 1:2, 1:5 and 1:10. After 24 hrs, the supernatant was removed, and cells were washed twice with 1x PBS before infecting with ZIKV (Mex 1-7 strain) [82] at an MOI of 1:0.1. After 4 days, supernatant was collected and ZIKV was quantified by focus forming assay [82]. The experiment was repeated three times.

### Gnotobiotic rearing and *in vivo* invasion in mosquitoes

*Ae. aegypti* gnotobiotic larvae were generated as previously described [83]. To synchronize hatching, sterile eggs were transferred to a conical flask and placed under a vacuum for 45 min. To verify sterility, larval water was plated on non-selective LB agar plates. Twenty L1 larvae were transferred to a T75 tissue culture flask and inoculated with transgenic symbionts possessing the pRAM18dRGA-mCherry at 1×10^7^. Bacterial cultures were quantified with a spectrophotometer (DeNovix DS-11, DeNovix) and validated by plating and determining colony forming units. L1 larvae grown without bacteria were used as contamination control, and these mosquitoes did not reach pupation [83]. To feed mosquitoes, ground fish food pellets were sterilized by autoclaving and then mixed with sterile water. 60 µl of fish food (1 µg/µl) was fed to larvae on alternative days.

## Acknowledgements

We would like to thank the UTMB insectary core for providing the lab mosquitoes. GLH is supported by NIH grants (R21AI138074, R21AI124452 and R21AI129507), the Wolfson Foundation and Royal Society (RSWF\R1\180013), the John S. Dunn Foundation Collaborative Research Award, and the Centers for Disease Control and Prevention (Cooperative Agreement Number U01CK000512). The paper contents are solely the responsibility of the authors and do not necessarily represent the official views of the Centers for Disease Control and Prevention or the Department of Health and Human Services. This work was also supported by a James W. McLaughlin postdoctoral fellowship at the University of Texas Medical Branch to SH and a NIH T32 fellowship (2T32AI007526) to MAS and Anti-VeC AV/PP0021//1 to AAS. Microscopy core facility at NYBC was supported by NYBC intramural fund. Confocal imaging facilities were funded by a Wellcome Trust Multi-User Equipment Grant (104936/Z/14/Z).

## Author Contributions

SH and GLH designed the experiments. SH, DV, ACS, MAS, and VLP completed the experiments. SH, DV, VLP, AKC, and GLH undertook analysis. SH, AKC, AAS, and GLH wrote and edited the manuscript and all authors agreed to the final version. GLH acquired the funding and supervised the work.

## Supplementary figure legends

**Figure S1. Gentamicin invasion assay in different cell lines.** Invasion of *E. coli* and *E. coli BL21* expressing the *Yersina invasion* gene (*Ypinv*) in different cell lines. The assay was done in Aag2 (*Aedes aegypti*), Sua5B (*Anopheles gambiae*) and Vero (Monkey kidney cells). The assay was done twice. The statistical significance was determined using an Unpaired t-test.

**Figure S2. Flourscent microscopy of bacteria in Aag2 cells.** Merged and separate channels – blue (DAPI), green (actin filaments stained with Phalloidin), red (bacteria expressing mCherry). Scale bars are 70 μm.

**Figure S3. Effect of intracellular bacteria on the cell viability**. Aag2 cell numbers at different times post invasion with *C. neteri.* Cells were supplement with gentamicin (200 µg/ml) or cultured in the absence of antibiotic.

**Figure S4. Validation of gene silencing in cells.** RT-qPCR analysis of beta (A) and alpha (B) integrin gene expression 24 hours post transfection of dsRNA. dsRNAs targetting GFP were used as the negative control. The experiment was repeated twice. The statistical significance was determined using an Unpaired t-test.

**Figure S5. CLSM analyses of infected adult gut and Malpighian tubule**. Maximum intensity projection of tissues 3D-reconstructed from a series of Z-stacks merging mosquito actin (white), mCherry-expressing *Cedecea* (yellow) and DAPI-stained DNA (magenta). Several XZ and YZ orthogonal views (OV) of the stacks are shown for each tissue on the sides. Scale bars are 50 μm.

**Figure S6.** Phylogenetic analysis of multilocus sequence typing as described in [76] shows the phylogenetic position of *Serratia sp.* Alb1 (in red; this study) and *C. neteri* Alb1 (in blue; [11]). The tree analysis was performed using PhyML under the general time-reversible model with rapid aLRT bootstrap support as implemented as default in seaview (v4.7; [77]. The *Serratia* sp. Alb1 sequences were submitted to NCBI under accession numbers XXX (ropB), XXX (gyrB), XXX (atpD) and XXX (infB). [Accession numbers requested and will be updated in revision process as soon as available].

## Supporting files

**Supporting file S1**. Tree file, including all support values.

## Supplementary table legends

**Table S1.** Primer sequences used in this study

## Supplementary video legends.

**Video S1**. Series of Z-stacks of an adult infected gut by CLSM.

**Video S2**. Series of Z-stacks of an adult infected Malpighian tubule by CLSM.

## Reference

1. Duguma D, Hall MW, Smartt CT, Neufeld JD. Effects of Organic Amendments on Microbiota Associated with the Culex nigripalpus Mosquito Vector of the Saint Louis Encephalitis and West Nile Viruses. mSphere. 2017;2(1). Epub 2017/02/09. doi:10.1128/mSphere.00387-16.

2. Dada N, Jumas-Bilak E, Manguin S, Seidu R, Stenström T-A, Overgaard HJ. Comparative assessment of the bacterial communities associated with Aedes aegypti larvae and water from domestic water storage containers. Parasites & Vectors. 2014;7(1):391. doi:10.1186/1756-3305-7-391.

3. Gusmao DS, Santos AV, Marini DC, Russo Ede S, Peixoto AM, Bacci Junior M, et al. First isolation of microorganisms from the gut diverticulum of Aedes aegypti (Diptera:Culicidae):new perspectives for an insect-bacteria association. Mem Inst Oswaldo Cruz. 2007;102(8):919–24. Epub 2008/01/23. doi:10.1590/s0074-02762007000800005.

4. Chavshin AR, Oshaghi MA, Vatandoost H, Yakhchali B, Zarenejad F, Terenius O. Malpighian tubules are important determinants of Pseudomonas transstadial transmission and longtime persistence in Anopheles stephensi. Parasites & Vectors. 2015;8(1):36. doi:10.1186/s13071-015-0635-6.

5. Chen S, Bagdasarian M, Walker ED. *Elizabethkingia anophelis*:Molecular Manipulation and Interactions with Mosquito Hosts. Applied and Environmental Microbiology. 2015;81(6):2233. doi:10.1128/AEM.03733-14.

6. Alonso DP, Mancini MV, Damiani C, Cappelli A, Ricci I, Alvarez MVN, et al. Genome Reduction in the Mosquito Symbiont Asaia. Genome Biol Evol. 2019;11(1):1–10. Epub 2018/11/27. doi:10.1093/gbe/evy255.

7. Duguma D, Hall MW, Smartt CT, Debboun M, Neufeld JD. Microbiota variations in Culex nigripalpus disease vector mosquito of West Nile virus and Saint Louis Encephalitis from different geographic origins. PeerJ. 2019;6:e6168. Epub 2019/01/16. doi:10.7717/peerj.6168.

8. Osei-Poku J, Mbogo CM, Palmer WJ, Jiggins FM. Deep sequencing reveals extensive variation in the gut microbiota of wild mosquitoes from Kenya. Molecular Ecology. 2012;21(20):5138–50. doi:10.1111/j.1365-294X.2012.05759.x.

9. Muturi EJ, Ramirez JL, Rooney AP, Kim C-H. Comparative analysis of gut microbiota of mosquito communities in central Illinois. PLoS Neglected Tropical Diseases. 2017;11(2):e0005377–18. doi:10.1371/journal.pntd.0005377.

10. Pei D, Jiang J, Yu W, Kukutla P, Uentillie A, Xu J. The waaL gene mutation compromised the inhabitation of Enterobacter sp. Ag1 in the mosquito gut environment. Parasites & Vectors. 2015:1–10. doi:10.1186/s13071-015-1049-1.

11. Hegde S, Nilyanimit P, Kozlova E, Anderson ER, Narra HP, Sahni SK, et al. CRISPR/Cas9-mediated gene deletion of the ompA gene in symbiotic Cedecea neteri impairs biofilm formation and reduces gut colonization of Aedes aegypti mosquitoes. PLOS Neglected Tropical Diseases. 2019;13(12):e0007883. doi:10.1371/journal.pntd.0007883.

12. Guegan M, Zouache K, Demichel C, Minard G, Tran Van V, Potier P, et al. The mosquito holobiont:fresh insight into mosquito-microbiota interactions. Microbiome. 2018;6(1):49. Epub 2018/03/21. doi:10.1186/s40168-018-0435-2.

13. Brady OJ, Godfray HC, Tatem AJ, Gething PW, Cohen JM, McKenzie FE, et al. Vectorial capacity and vector control:reconsidering sensitivity to parameters for malaria elimination. Trans R Soc Trop Med Hyg. 2016;110(2):107–17. Epub 2016/01/30. doi:10.1093/trstmh/trv113.

14. Hegde S, Rasgon JL, Hughes GL. The microbiome modulates arbovirus transmissionin mosquitoes. Current Opinion in Virology. 2015;15:97–102. doi:10.1016/j.coviro.2015.08.011.

15. Caragata EP, Tikhe CV, Dimopoulos G. Curious entanglements:interactions between mosquitoes, their microbiota, and arboviruses. Current Opinion in Virology. 2019;37:26–36. doi:https://doi.org/10.1016/j.coviro.2019.05.005.

16. Saldaña MA, Hegde S, Hughes GL. Microbial control of arthropod-borne disease. Memórias do Instituto Oswaldo Cruz. 2017;112(2):81–93. doi:10.1590/0074-02760160373.

17. Ricci I, Damiani C, Capone A, DeFreece C, Rossi P, Favia G. Mosquito/microbiota interactions:from complex relationships to biotechnological perspectives. Current Opinion in Microbiology. 2012;15(3):278–84. doi:10.1016/j.mib.2012.03.004.

18. Berhanu A, Abera A, Nega D, Mekasha S, Fentaw S, Assefa A, et al. Isolation and identification of microflora from the midgut and salivary glands of Anopheles species in malaria endemic areas of Ethiopia. BMC Microbiol. 2019;19(1):85. Epub 2019/05/01. doi:10.1186/s12866-019-1456-0.

19. Dickson LB, Ghozlane A, Volant S, Bouchier C, Ma L, Vega-Rua A, et al. Diverse laboratory colonies of Aedes aegypti harbor the same adult midgut bacterial microbiome. Parasites & Vectors. 2018;11(1):207. doi:10.1186/s13071-018-2780-1.

20. Damiani C, Ricci I, Crotti E, Rossi P, Rizzi A, Scuppa P, et al. Paternal transmission of symbiotic bacteria in malaria vectors. Current biology:CB. 2008;18(23):R1087–8. doi:10.1016/j.cub.2008.10.040.

21. Sharma P, Sharma S, Maurya RK, Das De T, Thomas T, Lata S, et al. Salivary glands harbor more diverse microbial communities than gut in Anopheles culicifacies. Parasites & Vectors. 2014;7(1):235. doi:10.1186/1756-3305-7-235.

22. Gimonneau G, Tchioffo MT, Abate L, Boissière A, Awono-Ambene PH, Nsango SE, et al. Composition of Anopheles coluzzii and Anopheles gambiae microbiota from larval to adult stages. Infection, genetics and evolution:journal of molecular epidemiology and evolutionary genetics in infectious diseases. 2014;28:715–24. doi:10.1016/j.meegid.2014.09.029.

23. Mancini MV, Damiani C, Accoti A, Tallarita M, Nunzi E, Cappelli A, et al. Estimating bacteria diversity in different organs of nine species of mosquito by next generation sequencing. BMC Microbiol. 2018;18(1):126. Epub 2018/10/06. doi:10.1186/s12866-018-1266-9.

24. Segata N, Baldini F, Pompon J, Garrett WS, Truong DT, Dabiré RK, et al. The reproductive tracts of two malaria vectors are populated by a core microbiome and by gender- and swarm-enriched microbial biomarkers. Scientific Reports. 2016;6:24207. doi:10.1038/srep24207.

25. Tchioffo MT, Boissiere A, Abate L, Nsango SE, Bayibeki AN, Awono-Ambene PH, et al. Dynamics of Bacterial Community Composition in the Malaria Mosquito’s Epithelia. Front Microbiol. 2015;6:1500. Epub 2016/01/19. doi:10.3389/fmicb.2015.01500.

26. Koosha M, Vatandoost H, Karimian F, Choubdar N, Oshaghi MA. Delivery of a Genetically Marked Serratia AS1 to Medically Important Arthropods for Use in RNAi and Paratransgenic Control Strategies. Microb Ecol. 2018. Epub 2018/11/22. doi:10.1007/s00248-018-1289-7.

27. Wang S, Dos-Santos ALA, Huang W, Liu KC, Oshaghi MA, Wei G, et al. Driving mosquito refractoriness to *Plasmodium falciparum* with engineered symbiotic bacteria. Science (New York, NY). 2017;357(6358):1399–402. doi:10.1126/science.aan5478.

28. Damiani C, Ricci I, Crotti E, Rossi P, Rizzi A, Scuppa P, et al. Mosquito-Bacteria Symbiosis:The Case of Anopheles gambiae and Asaia. Microb Ecol. 2010;60(3):644–54. doi:10.1007/s00248-010-9704-8.

29. Wei G, Lai Y, Wang G, Chen H, Li F, Wang S. Insect pathogenic fungus interacts with the gut microbiota to accelerate mosquito mortality. Proceedings of the National Academy of Sciences of the United States of America. 2017;114(23):5994–9. doi:10.1073/pnas.1703546114.

30. Nehme NT, Liégeois S, Kele B, Giammarinaro P, Pradel E, Hoffmann JA, et al. A model of bacterial intestinal infections in Drosophila melanogaster. 2007;3(11):e173. doi:10.1371/journal.ppat.0030173.

31. Czuczman MA, Fattouh R, van Rijn JM, Canadien V, Osborne S, Muise AM, et al. Listeria monocytogenes exploits efferocytosis to promote cell-to-cell spread. Nature. 2014;509(7499):230–4. Epub 2014/04/18. doi:10.1038/nature13168.

32. Ribet D, Cossart P. How bacterial pathogens colonize their hosts and invade deeper tissues. Microbes and infection. 2015;17(3):173–83. doi:10.1016/j.micinf.2015.01.004.

33. Hegde S, Hegde S, Spergser J, Brunthaler R, Rosengarten R, Chopra-Dewasthaly R. In vitro and in vivo cell invasion and systemic spreading of Mycoplasma agalactiae in the sheep infection model. International journal of medical microbiology:IJMM. 2014;304(8):1024–31. doi:10.1016/j.ijmm.2014.07.011.

34. Tchioffo MT, Abate L, Boissière A, Nsango SE, Gimonneau G, Berry A, et al. An epidemiologically successful Escherichia coli sequence type modulates Plasmodium falciparum infection in the mosquito midgut. Infection, genetics and evolution:journal of molecular epidemiology and evolutionary genetics in infectious diseases. 2016;43:22–30. doi:10.1016/j.meegid.2016.05.002.

35. Elsinghorst EA. Measurement of invasion by gentamicin resistance. Methods Enzymol. 1994;236:405–20. Epub 1994/01/01.

36. Isberg RR, Falkow S. A single genetic locus encoded by Yersinia pseudotuberculosis permits invasion of cultured animal cells by Escherichia coli K-12. Nature. 1985;317(6034):262–4.

37. Isberg RR, Voorhis DL, Falkow S. Identification of invasin:a protein that allows enteric bacteria to penetrate cultured mammalian cells. Cell. 1987;50(5):769–78. Epub 1987/08/28. doi:10.1016/0092-8674(87)90335-7.

38. Dersch P, Isberg RR. A region of the Yersinia pseudotuberculosis invasin protein enhances integrin-mediated uptake into mammalian cells and promotes self-association. The EMBO Journal. 1999;18(5):1199–213. doi:10.1093/emboj/18.5.1199.

39. Trujillo-Ocampo A, Cazares-Raga FE, Del Angel RM, Medina-Ramirez F, Santos-Argumedo L, Rodriguez MH, et al. Participation of 14-3-3epsilon and 14-3-3zeta proteins in the phagocytosis, component of cellular immune response, in Aedes mosquito cell lines. Parasit Vectors. 2017;10(1):362. Epub 2017/08/03. doi:10.1186/s13071-017-2267-5.

40. Dimopoulos G, Müller HM, Levashina EA, Kafatos FC. Innate immune defense against malaria infection in the mosquito. Current Opinion in Immunology. 2001;13(1):79–88.

41. Hegde S, Khanipov K, Albayrak L, Golovko G, Pimenova M, Saldaña MA, et al. Microbiome interaction networks and community structure from laboratory-reared and field-collected *Aedes aegypti*, *Aedes albopictus*, and *Culex quinquefasciatus* mosquito vectors. Frontiers in Microbiology. 2018;9:715. doi:10.3389/fmicb.2018.02160.

42. Gupta L, Molina-Cruz A, Kumar S, Rodrigues J, Dixit R, Zamora RE, et al. The STAT pathway mediates late-phase immunity against Plasmodium in the mosquito Anopheles gambiae. Cell Host & Microbe. 2009;5(5):498–507. doi:10.1016/j.chom.2009.04.003.

43. Wright JD, Barr AR. The ultrastructure and symbiotic relationships of Wolbachia of mosquitoes of the Aedes scutellaris group. Journal of Ultrastructure Research. 1980;72(1):52–64. doi:https://doi.org/10.1016/S0022-5320(80)90135-5.

44. Fattouh N, Cazevieille C, Landmann F. Wolbachia endosymbionts subvert the endoplasmic reticulum to acquire host membranes without triggering ER stress. PLOS Neglected Tropical Diseases. 2019;13(3):e0007218. doi:10.1371/journal.pntd.0007218.

45. Traven A, Naderer T. Microbial egress:a hitchhiker guide to freedom. PLOS Pathogens. 2014;10(7):e1004201. doi:10.1371/journal.ppat.1004201.

46. Flieger A, Frischknecht F, Hacker G, Hornef MW, Pradel G. Pathways of host cell exit by intracellular pathogens. Microb Cell. 2018;5(12):525–44. Epub 2018/12/12. doi:10.15698/mic2018.12.659.

47. Haglund CM, Welch MD. Pathogens and polymers:microbe-host interactions illuminate the cytoskeleton. J Cell Biol. 2011;195(1):7–17. Epub 2011/10/05. doi:10.1083/jcb.201103148.

48. Hybiske K, Stephens R. Cellular Exit Strategies of Intracellular Bacteria. Microbiol Spectr. 2015;3(6). Epub 2016/06/24. doi:10.1128/microbiolspec.VMBF-0002-2014.

49. Fukumatsu M, Ogawa M, Arakawa S, Suzuki M, Nakayama K, Shimizu S, et al. Shigella targets epithelial tricellular junctions and uses a noncanonical clathrin-dependent endocytic pathway to spread between cells. Cell Host & Microbe. 2012;11(4):325–36. doi:10.1016/j.chom.2012.03.001.

50. Casella JF, Flanagan MD, Lin S. Cytochalasin D inhibits actin polymerization and induces depolymerization of actin filaments formed during platelet shape change. Nature. 1981;293(5830):302–5. Epub 1981/09/24. doi:10.1038/293302a0.

51. Mizutani T, Kobayashi M, Eshita Y, Shirato K, Kimura T, Ako Y, et al. Involvement of the JNK-like protein of the Aedes albopictus mosquito cell line, C6/36, in phagocytosis, endocytosis and infection of West Nile virus. Insect Mol Biol. 2003;12. doi:10.1046/j.1365-2583.2003.00435.x.

52. Lin M, Kikuchi T, Brewer HM, Norbeck AD, Rikihisa Y. Global Proteomic Analysis of Two Tick-Borne Emerging Zoonotic Agents:Anaplasma Phagocytophilum and Ehrlichia Chaffeensis. Frontiers in Microbiology. 2011;2. doi:10.3389/fmicb.2011.00024.

53. Martinez JJ, Cossart P. Early signaling events involved in the entry of Rickettsia conorii into mammalian cells. Journal Of Cell Science. 2004;117(Pt 21):5097–106. doi:10.1242/jcs.01382.

54. Wesolowski J, Paumet F. Taking control:reorganization of the host cytoskeleton byChlamydia. F1000Research. 2017;6:2058. doi:10.12688/f1000research.12316.1.

55. Moita L, Vriend G, Mahairaki V, Louis C, Kafatos F. Integrins of Anopheles gambiae and a putative role of a new β integrin, BINT2, in phagocytosis of E. coli. Insect β Biochemistry and Molecular Biology. 2006;36(4):282–90. doi:10.1016/j.ibmb.2006.01.004.

56. Eto DS, Jones TA, Sundsbak JL, Mulvey MA. Integrin-mediated host cell invasion by type 1-piliated uropathogenic Escherichia coli. PLOS Pathogens. 2007;3(7):e100. doi:10.1371/journal.ppat.0030100.

57. Wu J, Weening EH, Faske JB, Höök M, Skare JT. Invasion of Eukaryotic Cells by 1Integrins and Src Kinase Activity. Infection and β Immunity. 2011;79(3):1338–48. doi:10.1128/IAI.01188-10.

58. Gillenius E, Urban CF. The adhesive protein invasin of Yersinia pseudotuberculosis induces neutrophil extracellular traps via β ntegrins. Microbes and infection. 2015;17(5):327–36. doi:10.1016/j.micinf.2014.12.014.

59. Miesen P, van Rij RP. Crossing the Mucosal Barrier:A Commensal Bacterium Gives Dengue Virus a Leg-Up in the Mosquito Midgut. Cell Host Microbe. 2019;25(1):1–2. Epub 2019/01/11. doi:10.1016/j.chom.2018.12.009.

60. Coatsworth H, Caicedo PA, Van Rossum T, Ocampo CB, Lowenberger C. The Composition of Midgut Bacteria in Aedes aegypti (Diptera:Culicidae) That Are Naturally Susceptible or Refractory to Dengue Viruses. Journal of Insect Science. 2018;18(6). doi:10.1093/jisesa/iey118.

61. Hegde S, Hegde S, Hegde S, Zimmermann M, Flöck M, Spergser J, et al. Simultaneous Identification of Potential Pathogenicity Factors of Mycoplasma agalactiae in the Natural Ovine Host by Negative Selection. Infection and Immunity. 2015;83(7):2751–61. doi:10.1128/IAI.00403-15.

62. Broderick NA. Friend, foe or food? Recognition and the role of antimicrobial peptides in gut immunity and Drosophila-microbe interactions. Philosophical transactions of the Royal Society of London Series B, Biological sciences. 2016;371(1695):20150295. doi:10.1098/rstb.2015.0295.

63. Buchon N, Broderick NA, Poidevin M, Pradervand S, Lemaitre B. Drosophila intestinal response to bacterial infection:activation of host defense and stem cell proliferation. Cell Host & Microbe. 2009;5(2):200–11. doi:10.1016/j.chom.2009.01.003.

64. Cheng G, Liu Y, Wang P, Xiao X. Mosquito Defense Strategies against Viral Infection. Trends Parasitol. 2016;32(3):177–86. Epub 2015/12/03. doi:10.1016/j.pt.2015.09.009.

65. Fragkoudis R, Attarzadeh-Yazdi G, Nash AA, Fazakerley JK, Kohl A. Advances in dissecting mosquito innate immune responses to arbovirus infection. The Journal of general virology. 2009;90(Pt 9):2061–72. doi:10.1099/vir.0.013201-0.

66. Zhang X, Aksoy E, Girke T, Raikhel AS, Karginov FV. Transcriptome-wide microRNA and target dynamics in the fat body during the gonadotrophic cycle of Aedes aegypti. Proceedings of the National Academy of Sciences of the United States of America. 2017;114(10):E1895–E903. doi:10.1073/pnas.1701474114.

67. Barletta AB, Nascimento-Silva MC, Talyuli OA, Oliveira JH, Pereira LO, Oliveira PL, et al. Microbiota activates IMD pathway and limits Sindbis infection in Aedes aegypti. Parasit Vectors. 2017;10(1):103. Epub 2017/02/25. doi:10.1186/s13071-017-2040-9.

68. Kumar A, Srivastava P, Sirisena P, Dubey SK, Kumar R, Shrinet J, et al. Mosquito Innate Immunity. Insects. 2018;9(3). Epub 2018/08/12. doi:10.3390/insects9030095.

69. Friedrich N, Hagedorn M, Soldati-Favre D, Soldati T. Prison break:pathogens strategies to egress from host cells. Microbiol Mol Biol Rev. 2012;76(4):707–20. doi:10.1128/MMBR.00024-12.

70. Narasimhan S, Rajeevan N, Liu L, Zhao YO, Heisig J, Pan J, et al. Gut Microbiota of the Tick Vector Ixodes scapularis Modulate Colonization of the Lyme Disease Spirochete. Cell Host & Microbe. 2014;15(1):58–71. doi:10.1016/j.chom.2013.12.001.

71. Rodgers FH, Gendrin M, Wyer CAS, Christophides GK. Microbiota-induced peritrophic matrix regulates midgut homeostasis and prevents systemic infection of malaria vector mosquitoes. PLOS Pathogens. 2017;13(5):e1006391. doi:10.1371/journal.ppat.1006391.

72. Gusmão DS, Santos AV, Marini DC, Bacci Jr M, Berbert-Molina MA, Lemos FJA. Culture-dependent and culture-independent characterization of microorganisms associated with Aedes aegypti (Diptera:Culicidae) (L.) and dynamics of bacterial colonization in the midgut. Acta Tropica. 2010;115(3):275–81. doi:10.1016/j.actatropica.2010.04.011.

73. Tsai YL, Hayward RE, Langer RC, Fidock DA, Vinetz JM. Disruption of Plasmodium falciparum Chitinase Markedly Impairs Parasite Invasion of Mosquito Midgut. Infection and Immunity. 2001;69(6):4048–54. doi:10.1128/IAI.69.6.4048-4054.2001.

74. Goebel W, Gross R. Intracellular survival strategies of mutualistic and parasitic prokaryotes. Trends Microbiol. 2001;9(6):267–73. Epub 2001/06/08.

75. Kumar S, Molina-Cruz A, Gupta L, Rodrigues J, Barillas-Mury C. A peroxidase/dual oxidase system modulates midgut epithelial immunity in *Anopheles gambiae*. Science. 2010;327(5973):1644–8. doi:10.1126/science.1184008.

76. Brady C, Cleenwerck I, Venter S, Coutinho T, De Vos P. Taxonomic evaluation of the genus Enterobacter based on multilocus sequence analysis (MLSA):Proposal to reclassify E. nimipressuralis and E. amnigenus into Lelliottia gen. nov. as Lelliottia nimipressuralis comb. nov. and Lelliottia amnigena comb. nov., respectively, E. gergoviae and E. pyrinus into Pluralibacter gen. nov. as Pluralibacter gergoviae comb. nov. and Pluralibacter pyrinus comb. nov., respectively, E. cowanii, E. radicincitans, E. oryzae and E. arachidis into Kosakonia gen. nov. as Kosakonia cowanii comb. nov., Kosakonia radicincitans comb. nov., Kosakonia oryzae comb. nov. and Kosakonia arachidis comb. nov., respectively, and E. turicensis, E. helveticus and E. pulveris into Cronobacter as Cronobacter zurichensis nom. nov., Cronobacter helveticus comb. nov. and Cronobacter pulveris comb. nov., respectively, and emended description of the genera Enterobacter and Cronobacter. Systematic and Applied Microbiology. 2013;36(5):309–19. doi:https://doi.org/10.1016/j.syapm.2013.03.005.

77. Gouy M, Guindon S, Gascuel O. SeaView version 4:A multiplatform graphical user interface for sequence alignment and phylogenetic tree building. Molecular biology and evolution. 2010;27(2):221–4. Epub 2009/10/23. doi:10.1093/molbev/msp259.

78. Peleg J. Growth of arboviruses in primary tissue culture of Aedes aegypti embryos. Am J Trop Med Hyg. 1968;17.

79. Horn T, Boutros M. E-RNAi:a web application for the multi-species design of RNAi reagents--2010 update. Nucleic Acids Research. 2010;38(Web Server issue):W332–9. doi:10.1093/nar/gkq317.

80. Livak KJ, Schmittgen TD. Analysis of Relative Gene Expression Data Using Real-Time Quantitative PCR and the 2−ΔΔCT Method. Methods (San Diego, Calif). 2001;25(4):402–8. doi:10.1006/meth.2001.1262.

81. Isoe J, Collins J, Badgandi H, Day WA, Miesfeld RL. Defects in coatomer protein I (COPI) transport cause blood feeding-induced mortality in Yellow Fever mosquitoes. Proceedings of the National Academy of Sciences of the United States of America. 2011;108(24):E211–7. doi:10.1073/pnas.1102637108.

82. Saldaña MA, Etebari K, Hart CE, Widen SG, Wood TG, Thangamani S, et al. Zika virus alters the microRNA expression profile and elicits an RNAi response in Aedes aegypti mosquitoes. PLoS Neglected Tropical Diseases. 2017;11(7):e0005760–18. doi:10.1371/journal.pntd.0005760.

83. Coon KL, Vogel KJ, Brown MR, Strand MR. Mosquitoes rely on their gut microbiota for development. Molecular Ecology. 2014;23(11):2727–39. doi:10.1111/mec.12771.

