## Supplementary material for "Gut-associated bacteria invade the midgut epithelium of *Aedes aegypti* and stimulate innate immunity and suppress Zika virus infection in cells": Table S1

| Primer | Primer sequnce (5'-3') | Reference |
| --- | --- | --- |
| QPCT_AAEL000709R | CGCTCCGGTAGCCTCGTGGATC | This study |
| QPCR_AAEL000709F | AGACAGCCGCACCTTCGATTC | This study |
| QPCR_AAEL0007696R | CTGCCTGCGTGACCGTATCC | This study |
| QPCT_AAEL007696F | TGGTGGTGGTGTCCTGCGTAAC | This study |
| qPCR_Caspar_F | TGTACGGGTGGGATCTAACA | This study |
| qPCR_Caspar_R | AGGAATGTTTCGGACCGTTATC | This study |
| Isoe aegypti S7 F | ACCGCCGTCTACGATGCCA | Isoe et al |
| Isoe aegypti S7 R | ATGGTGGTCTGCTGGTTCTT | Isoe et al |
| AeREL2_RTF | GGACGAGGCAGCGGCGCAGTTTGAGC | This study |
| AeREL2_RTR | TCCAGAGGGCCGAGATAAGTTCC | This study |
| AAEL004522_GAM_F | GCCAAAACCTGTTCCTCTTG | Telang et al |
| AAEL004522_GAM_R | CGATGTAGCATTCGGTGATG | Telang et al |
| AAEL003832_DEFC_F | TTGTTTGCTTCGTTGCTCTTT | Telang et al |
| AAEL003832_DEFC_R | ATCTCCTACACCGAACCCACT | Telang et al |
| AAEL015515_CECG_F | TCACAAAGTTATTTCTCCTGATCG | Telang et al |
| AAEL015515_CECG_R | GCTTTAGCCCCAGCTACAAC | Telang et al |
| 16S univR | CCATTGTAGCACGTGT | O'Neill et al. 1992 |
| 16S univF | GCTTAACACATGCAAG | O'Neill et al. 1992 |
| T7_Ae_betaint2_F | TAATACGACTCACTATAGGGCAGCACCGAGAGGGATGTATG | This study |
| T7_Ae_betaint2_R | TAATACGACTCACTATAGGGAATACCTGCATTAACGCGTCC | This study |
| Ae_betaint_RTF1 | TAGGCCACAGCGCCTTAAAA | This study |
| Ae_betaint_RTR1 | GGCTCACATACATCCCTCTCG | This study |
| Ae.alpha1T7_F1 | TAA TAC GAC TCA CTA TAG GGC TAC AGA GAT ACG TAC AGC G | This study |
| Ae.alpha1T7_R1 | TAA TAC GAC TCA CTA TAG GGC TTC ATT CAC CTG TCG CCT G | This study |
| Aealpha1_RTF | TGC GGT CCA AAT TGA ATG CG | This study |
| Aealpha1_RTR | GGG ACA AAA TCT GCA GGG CAA | This study |

Telang A, Qayum AA, Parker A, Sacchetta BR, Byrnes GR. Larval nutritional stress affects vector immune traits in adult yellow fever mosquito Aedes aegypti (Stegomyia aegypti). Medical and Veterinary Entomology. 2012;26(3):271-81. doi: 10.1111/j.1365-2915.2011.00993.x.

O’Neill SL, Giordano R, Colbert AM, Karr TL, Robertson HM. 16S rRNA phylogenetic analysis of the bacterial endosymbionts associated with cytoplasmic incompatibility in insects. Proceedings of the National Academy of Sciences of the United States of America. 1992;89(7):2699-702.

Isoe J, Collins J, Badgandi H, Day WA, Miesfeld RL. Defects in coatomer protein I (COPI) transport cause blood feeding-induced mortality in Yellow Fever mosquitoes. Proceedings of the National Academy of Sciences of the United States of America. 2011;108(24):E211-7. doi: 10.1073/pnas.1102637108.
