## Supplementary figures and images for "Gut-associated bacteria invade the midgut epithelium of *Aedes aegypti* and stimulate innate immunity and suppress Zika virus infection in cells"

### Figure S1

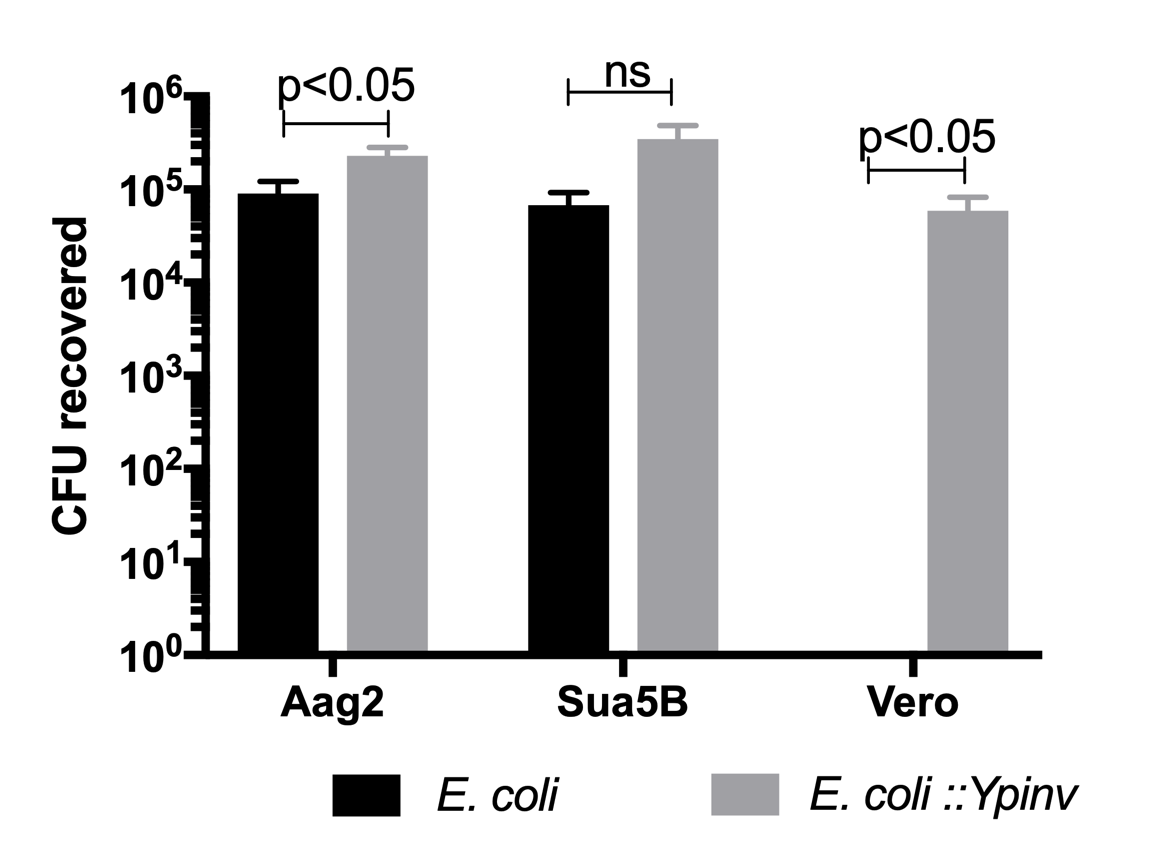

### Figure S2

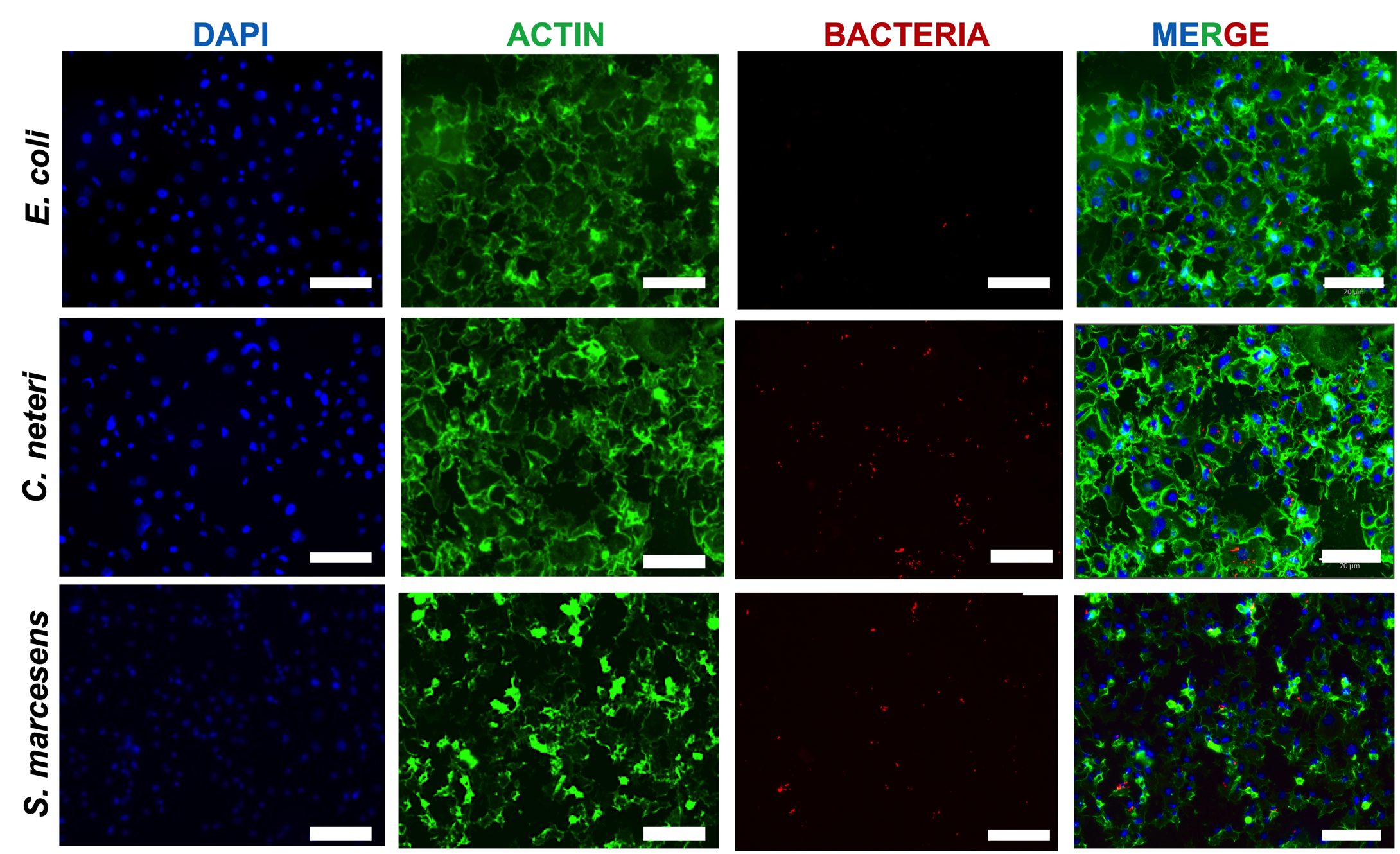

### Figure S3

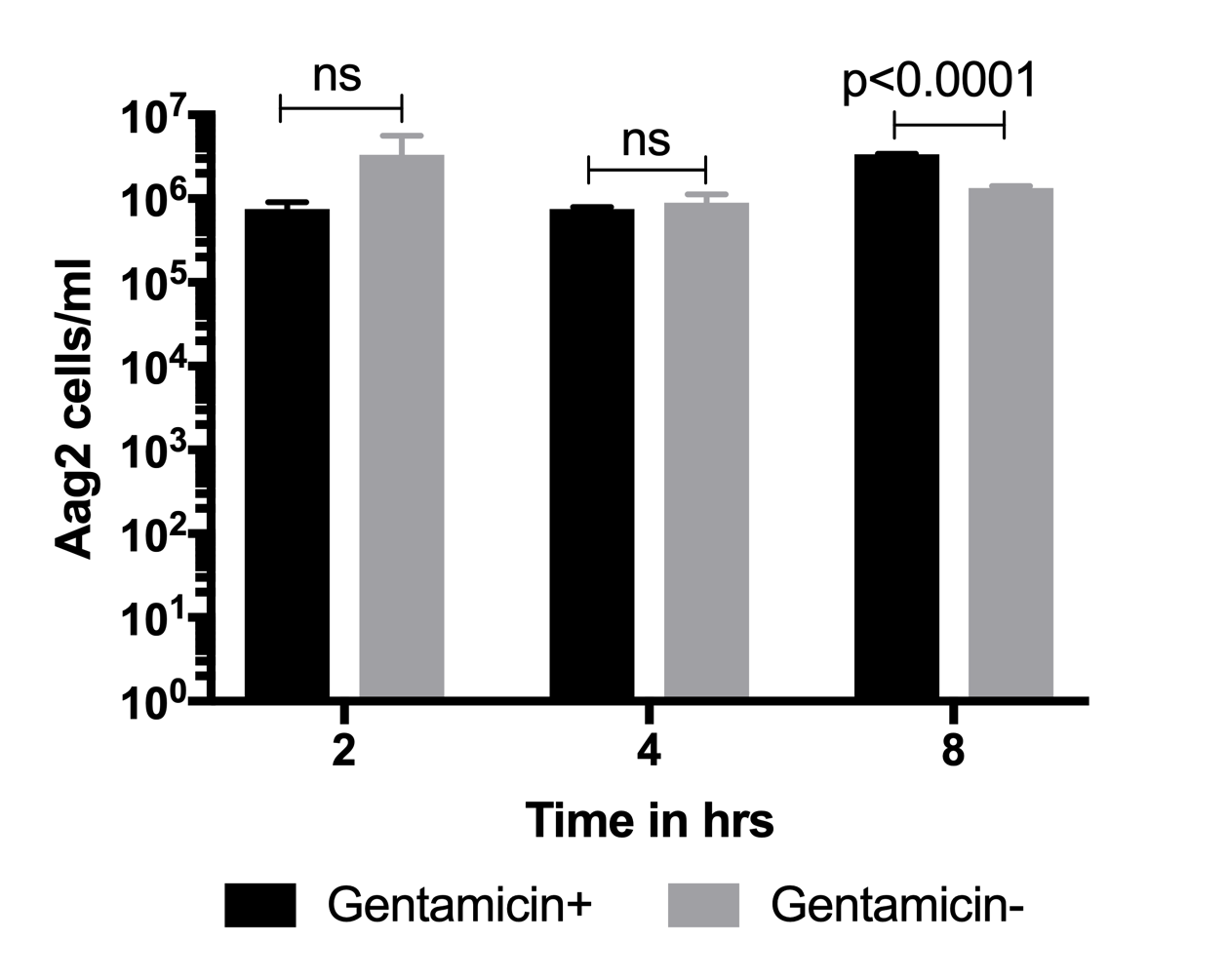

### Figure S4

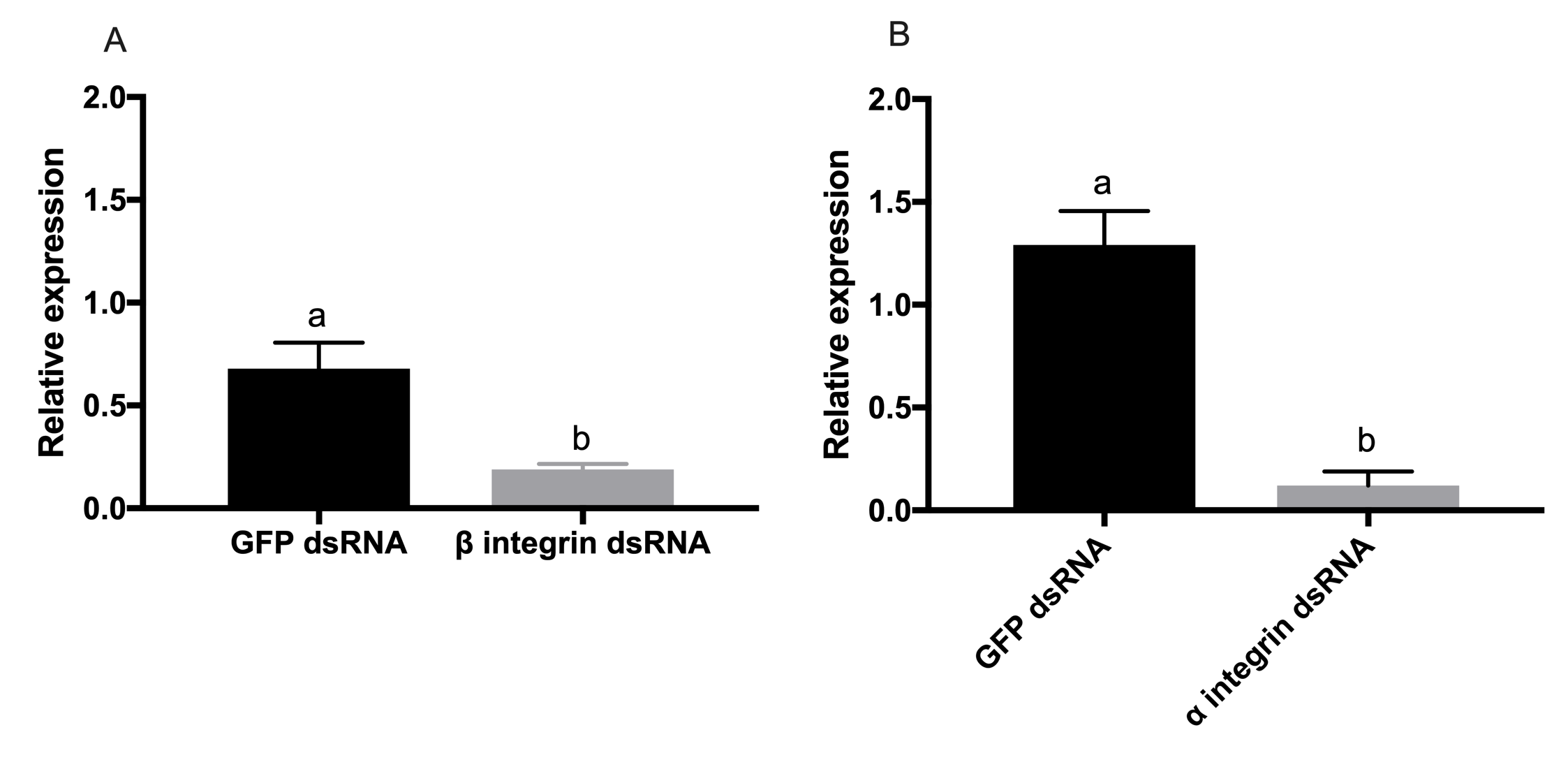

### Figure S5

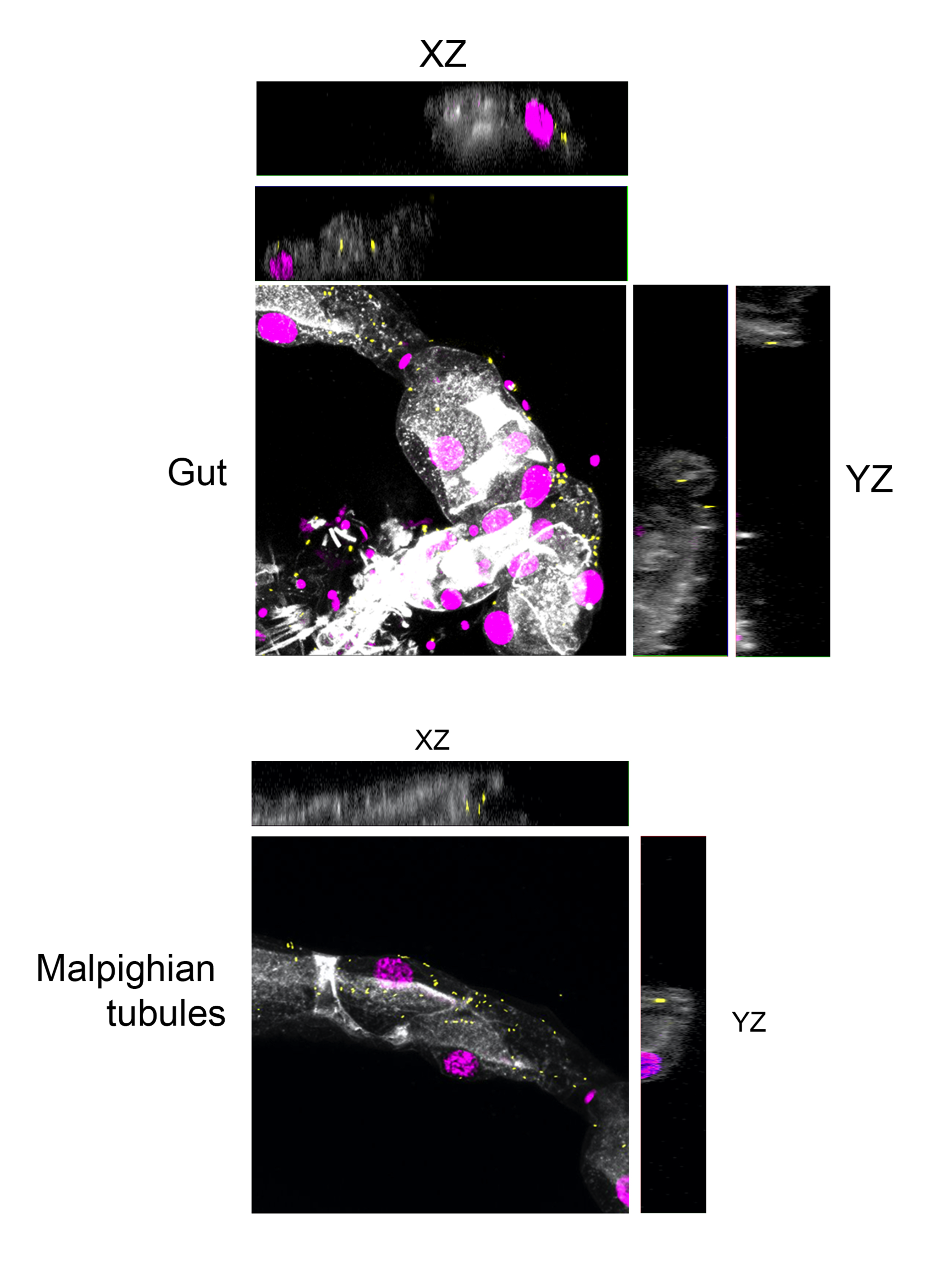

### Figure S6

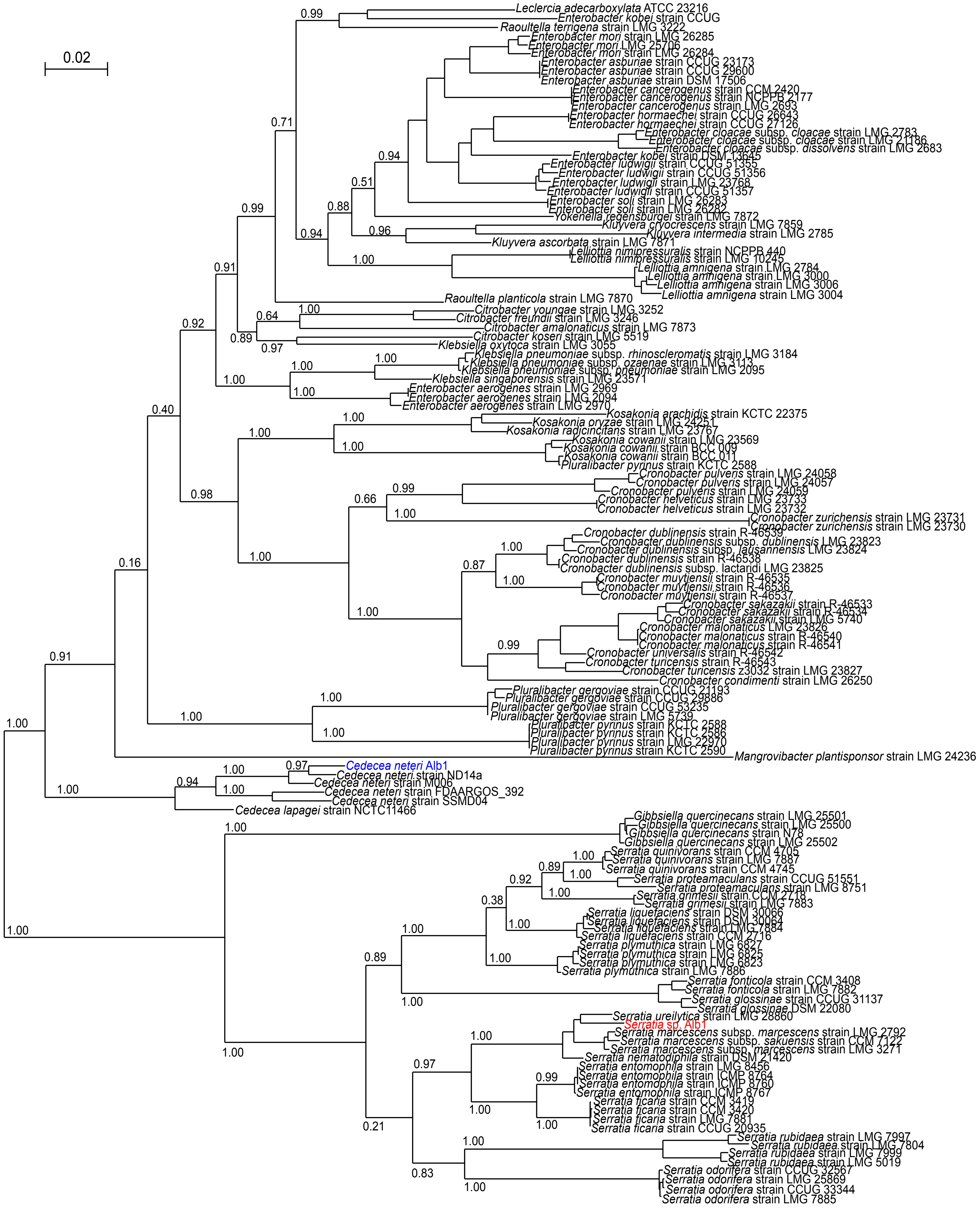
